# Atacformer: A transformer-based foundation model for analysis and interpretation of ATAC-seq data

**DOI:** 10.1101/2025.11.03.685753

**Authors:** Nathan J. LeRoy, Guangtao Zheng, Oleksandr Khoroshevskyi, Donald R Campbell, Aidong Zhang, Nathan C. Sheffield

## Abstract

**Introduction:** Chromatin accessibility profiling is an important tool for understanding gene regulation and cellular function. While public repositories house nearly 10,000 scATAC-seq experiments, unifying this data for meaningful analysis remains challenging. Existing tools struggle with the scale and complexity of scATAC-seq datasets, limiting tasks like clustering, cell-type annotation, and reference mapping. A promising solution is using foundation models adapted to specific tasks via transfer learning. While transfer learning has been applied to scRNA-seq, its potential for scATAC-seq remains underexplored.

**Methods:** We introduce Atacformer, a transformer-based foundation model for scATAC-seq data analysis. Unlike other models that only produce cell-level representations, Atacformer generates embeddings for individual cis-regulatory elements. Pre-trained on a large atlas of scATAC-seq experiments, Atacformer learns robust representations of genomic regulatory regions for downstream use. After pretraining, the model is fine-tuned for cell-type prediction and batch correction. We also integrated Atacformer with RNA-seq data to build a Contrastive RNA-ATAC Fine Tuning (CRAFT) model capable of cross-modal alignment and RNA imputation from ATAC data.

**Results:** Atacformer matches or exceeds leading scATAC-seq clustering tools in adjusted rand index and runtime, with fine-tuned models achieving top performance across datasets. It processes raw fragment files end-to-end 80% faster than existing tools while preserving biological structure. Fine-tuned on bulk BED files, it recovers cell type and assay labels with *>*80% accuracy. We show how the Atacformer architecture produces contextualized embeddings of individual genomic regions, which we use to identify unannotated, cell-type-specific promoter elements directly from chromatin accessibility data.

## Introduction

The exponential growth of publicly available genomic data has spurred the development of powerful pre-trained foundation models across diverse genomic modalities, including DNA sequences (*e.g*. Enformer^1^ and DNA Discrete Diffusion^2^); gene expression (*e.g*. Geneformer^3^ and scGPT^4^); and protein folding (*e.g*. AlphaFold^5–7^). By leveraging transfer learning, these models have dramatically enhanced data integration capabilities, enabling researchers to apply knowledge learned from large-scale datasets to new, often smaller-scale tasks.

Despite significant progress across genomics, transcriptomics, and proteomics, the development of foundation models explicitly tailored for epigenomic data remains comparatively underexplored. The Assay for Transposase-Accessible Chromatin sequencing (ATAC-seq) and its single-cell variant (scATAC-seq)^8,9^ have emerged as key methods for interrogating the regulatory landscape of the genome, offering insights into disease mechanisms, cellular heterogeneity, and tissue development. However, these assays present computational challenges due to their high dimensionality and inherent data sparsity^10,11^. Consequently, a wide array of computational tools have been developed to manage these complexities and simplify analysis pipelines for researchers. Comprehensive platforms, including ArchR^12^ and SnapATAC^13,14^, provide scalable, end-to-end solutions that support many standard workflows, such as quality control, dimensionality reduction, and trajectory analysis. More advanced, specialized tools like cisTopic provide a probabilistic topic modeling framework for discovering co-accessible enhancers and regulatory patterns, while deep learning approaches like SCALE^15^ and scBasset^16^ seek to address fundamental challenges of data sparsity and learning relationships between DNA sequence and chromatin accessibility.

These tools have pushed forward the analysis of scATAC-seq data. However, current methods for scATAC-seq are specialized, task-specific, and limited to the immediate dataset under investigation. They fail to exploit the shared biological knowledge encoded in the many publicly available datasets. No method has focused on producing a foundation model, trained on vast and diverse datasets, designed to be adapted to both new models through downstream fine-tuning and to new datasets through transfer learning. Recognizing this limitation, new models for scATAC-seq are emerging. This includes ChromFound^17^ and EpiAgent^18^, which are the first foundation models explicitly tailored for scATAC-seq, utilizing large-scale pre-training on millions of cells. Despite their advances, these models have several limitations. First, they are confined to single-cell scATAC-seq data and do not handle bulk ATAC-seq region-sets, which are widely used in many contexts. Second, they do not model genomic regions as discrete tokens, opting instead for continuous representations of cells that limit interpretability. Third, they rely on very large architectures – for example, EpiAgent uses over 1 billion parameters across its embedding and transformer modules, which requires substantial compute power for inference and adoption. Fourth, they do not explore multimodal contrastive integration with scRNA-seq, which restricts cell-type transfer and cross-modal inference capabilities^19^.

To address these limitations, we present Atacformer, a transformer-based foundation model. Atacformer leverages large-scale pre-training on single-cell ATAC-seq data. Furthermore, unlike existing approaches, we model genomic intervals as discrete tokens – the fundamental “words” of the regulatory genome. This token-based representation aligns with biological intuition and allows Atacformer to exploit the strengths of transformer architectures, which are particularly well-suited for learning from sequences of discrete inputs, as demonstrated in Natural Language Processing. Beyond the Atacformer model, we also introduce a dual-encoder contrastive learning approach called Contrastive RNA-ATAC-Fine-Tuning (CRAFT), which supports cross-modal alignment between scATAC-seq and scRNA-seq data. We benchmarked the performance of Atacformer and CRAFT models across four key applications. First, we assessed zero-shot cell clustering on multiple new, unseen PBMC datasets after fine-tuning the model on a cell-type clustering task. Next, we tested direct fragment file processing to measure speed and biological concordance without conventional preprocessing. Third, we applied Atacformer to bulk region-set data to evaluate embedding quality and metadata prediction. Finally, we demonstrate that Atacformer’s contextualized region embeddings can identify unannotated promoter regions that escape detection by conventional genome annotations but show cell-type-specific regulatory activity. Our results demonstrate that Atacformer can achieve state-of-the-art performance with best-in-class runtime for scATAC-seq analysis while maintaining powerful, zero-shot clustering performance on new, unseen datasets, and revealing hidden layers of cell-type-specific gene regulation through the discovery of functionally active but unannotated promoter elements.

## Results

### Atacformer is a new transformer-based foundation model for ATAC-seq data

Atacformer is a transformer-based^20^ foundation model that produces contextualized embeddings of genomic regions. Unlike existing models that are large, rigid, and task-specific, Atacformer is general-purpose and lightweight: users need only provide chromosome coordinates (chromosome, start, end) to begin analysis. By minimizing assumptions about the data and streamlining inputs, Atacformer serves as a fast, efficient backbone that can be adapted to many genomic interval tasks. Atacformer consists of two main components: a genomic region embedding module and a stacked transformer encoder layer with multi-head attention (Fig. 1A). To train Atacformer, we curated a single-cell ATAC-seq atlas consisting of 1.2 million cells and 10 billion tokens from 30 tissues (Fig. 1B; Table S1)^21–26^. We uniformly processed all raw datasets using a standardized pre-processing pipeline to ensure data integrity and compatibility (see Methods). We used these uniformly processed results to create a unified consensus vocabulary based on our earlier work^27^ consisting of 890,704 distinct genomic regions (see Methods).

**Figure 1.**
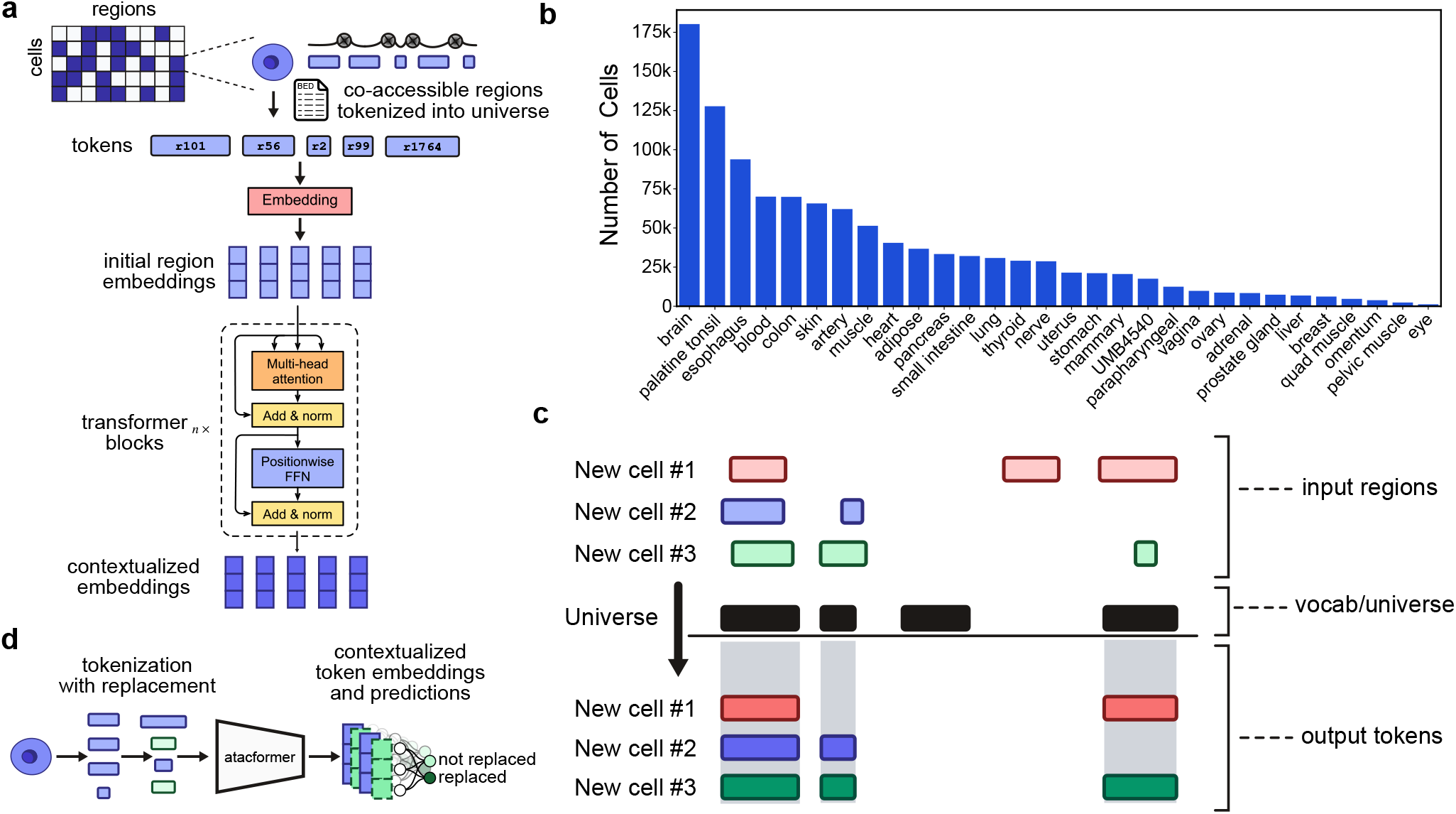
Atacformer architecture and overview of pretraining procedure. **a**. Model architecture and pretraining schematic for Atacformer. Individual cells are tokenized into the model universe, followed by random token replacement. These tokens are then passed to the embedding module, followed by n transformer blocks to generate contextualized embeddings. **b**. Tissue distribution and representation in the scATAC atlas used in Atacformers pretraining. **c**. The Atacformer tokenization strategy. New cells are tokenized into the model vocabulary using interval overlap analysis.

To tokenize a single cell into a set of dense, low-dimensional embeddings, we first map each accessible region in the cell to a corresponding region in the model’s vocabulary through simple interval intersection (Fig. 1C). This process transforms noisy, unstandardized genomic intervals into fixed tokens while preserving the biological significance of co-accessibility patterns. To enable extremely fast, in-memory tokenization that supports modern machine learning workflows, we developed a set of Rust-based tokenizers to be used in conjunction with Atacformer^28^.

Atacformer is trained using an ELECTRA-style pre-training objective^29^ (Fig. 1D) in which Atacformer receives tokenized region sets in which a random subset of tokens are replaced with others sampled from the vocabulary. The model is then tasked with predicting which tokens were replaced. Unless noted, we pre-trained Atacformer using a 45% token replacement rate and a context window of 8,192, as this captures the majority of co-accessible regions in all single-cells in our corpus (Fig. S1). We used a variety of training callbacks to determine the most optimal training time for our foundation model (Fig. S2). We refer to this model as `atacformer-base`. We evaluated `atacformer-base` to establish a performance baseline, and then fine-tuned it to achieve better performance and enable more flexible downstream analyses.

### Atacformer can be paired with Geneformer for powerful multiomics analysis

We sought to fine-tune Atacformer on a multimodal task. Recent breakthroughs, such as CLIP^30^, demonstrate that aligning fundamentally different data types in a shared latent space enables zero-shot transfer and cross-modal retrieval. Inspired by this, we investigated whether Atacformer would benefit from being paired with an encoder of another modality, such as scRNA-seq. To investigate, we developed Contrastive RNA-ATAC Fine-Tuning (CRAFT), a multi-modal model that combines Atacformer with Geneformer, a transcriptome encoder^3^. CRAFT uses contrastive training on multiomic data to create a shared latent space for scATAC-seq and scRNA-seq data. We initialized the model with pre-trained Atacformer and Geneformer models. We then trained CRAFT using a multiomic dataset containing over 106,000 cells profiled with scATAC and scRNA simultaneously (Fig. 2A; see Methods).

**Figure 2.**
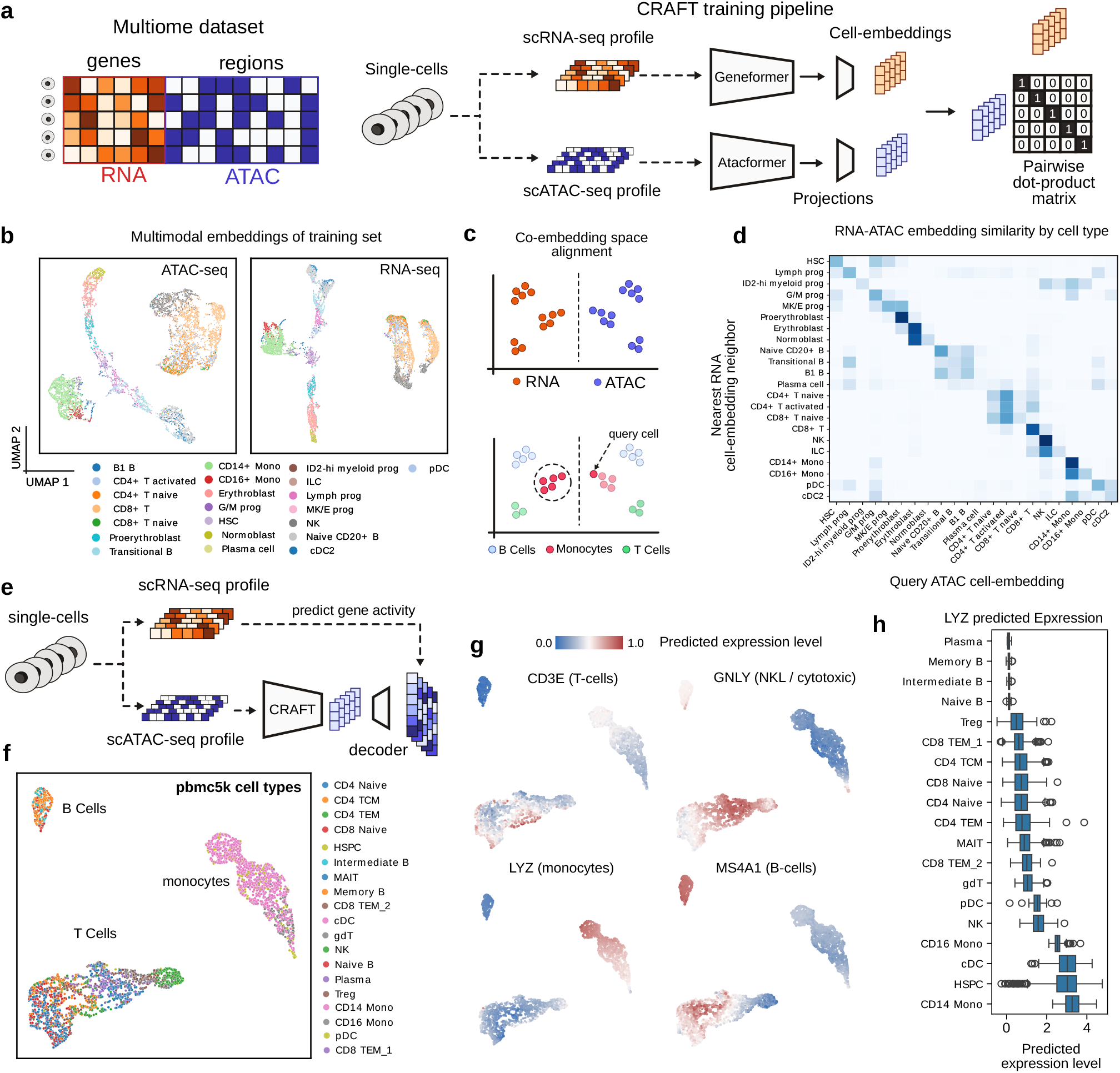
CRAFT is a powerful dual-encoder, multimodal single-cell embedding model. **a**. Schematic of the CRAFT training procedure. (Left) Schematic of a multiomic single-cell dataset showing ATAC and RNA signal jointly profiled in a single cell. (Right) Training step for a single mini-batch of cells. **b**. UMAP visualizations of the single-cell embeddings generated using the ATAC-encoder (left) and the RNA encoder (right). **c**. Resultant embedding space after training a CLIP-style multimodal model on single-cell RNA-seq and ATAC-seq data. (Top) Before training, RNA (orange) and ATAC (blue) measurements exist in separate feature spaces. (Bottom) The trained model projects both modalities into a shared embedding space where the same cell types (B cells, monocytes, T cells) co-localize regardless of assay modality. **d**. Heatmap of nearest RNA-embedding neighbors to a single ATAC-embedding cell, organized by cell type. **e**. Schematic of the RNA-decoder training procedure. **f**. PBMC5k dataset clustered using ATAC-embeddings; colored by cell-type. **g**. Heatmaps of PBMC5k dataset, colored by predicted RNA-expression profile for four different marker genes. **h**. Distribution of predicted LYZ expression levels across cell types.

UMAP visualizations of ATAC-seq and RNA-seq embeddings were topologically similar, highlighting modality alignment (Fig. 2B). Embeddings maintained biologically meaningful relationships; for example, the distinct groups of monocytes and CD8+ T cells are maintained in both modalities, as is a clear linear trajectory of the stages of red blood cell development from proerythroblast to erythroblast to normoblast. When projected into a single two-dimensional UMAP, modalities separated visually (Fig. S3); but despite this, nearest neighbors in higher-dimensional spaces preserved biological similarities, allowing accurate cross-modal cell-type alignment (Fig. 2C). To assess biological coherence, we projected ATAC embeddings into the multimodal CRAFT latent space and queried their nearest RNA neighbors. Consistently, the local RNA neighborhoods aligned with the same cell type as the corresponding ATAC embedding. (Fig. 2D).

Next, we assessed whether the aligned ATAC embeddings encoded transcriptomic information by training a small decoder to predict RNA-seq profiles from ATAC embeddings (Methods, Fig. 2E). This should allow us to leverage the joint CRAFT embeddings to predict scRNA-seq outputs from datasets where only scATAC-seq was assayed, or vice versa. We applied this decoder to an unseen scATAC-seq dataset with known labels, labeled with scVI (see methods). UMAP projections of the ATAC embeddings visually separated cells into clusters for T Cell, B Cell, and Monocyte lineages (Fig. 2F), showing that these out-of-sample cells are distinguished by the model. Next, we applied the RNA-seq decoder to predict scRNA-seq profiles. The predicted RNA-seq profiles accurately recapitulated known gene expression differences among these lineages, including cell-type-specific markers. For example, the imputed expression of monocyte marker LYZ was elevated in monocytes; B cell marker MS4A1 was elevated in B cells; T cell marker CD3E was elevated in T cells; cytotoxic cell marker GNLY was elevated in cytotoxic cells (Fig. 2G). Finally, we examined the predicted normalized expression across cell-types quantitatively. As an example, we show that the monocyte marker LYZ is disproportionately upregulated in Monocyte-like cells (Fig. 2H). Collectively, these results demonstrate that RNA-seq patterns have been incorporated in the ATAC-seq embeddings of the multi-modal CRAFT model, and that CRAFT integrates Atacformer embeddings with complementary single-cell models, enabling robust multimodal analysis and cross-modal biological inference. This dual-encoder framework accurately aligns chromatin accessibility and transcriptomic data within a unified latent space, facilitating precise cell-type identification and enhancing the interpretability of single-cell chromatin profiles.

### Fine-tuned Atacformer models and CRAFT enable fast and accurate zero-shot cell-clustering

The first bottleneck in scATAC-seq analysis is cell clustering. Most current tools require significant processing times (minutes per thousand cells), and many even require retraining on each dataset. We sought to make clustering faster while retaining or improving accuracy. A pre-trained Atacformer model could possibly cluster unseen data very quickly; however, we reasoned that a model trained on an unsupervised token replacement prediction task would perform sub-optimally for a cell-type clustering task. To that end, we sought to tune the base embeddings in two ways: First, inspired by the Sentence-BERT (SBERT) family of text embedding models^31^, we designed a novel fine-tuning approach for cell-type clustering. Starting with the base model (`atacformer-base`), we trained a *cell-type fine-tuned* model, `atacformer-ctft` using triplet loss to position cells of the same type together in the latent space (Fig. 3A; see Methods). We used the Luecken2021 multi-omics PBMC dataset for fine-tuning, which provides high-confidence cell-type labels across matched modalities^32^. Second, we also tested CRAFT, reasoning that CRAFT’s mutual refinement of ATAC-seq and RNA-seq signal could improve performance on downstream tasks,a phenomenon seen in other contexts^33,34^.

**Figure 3.**
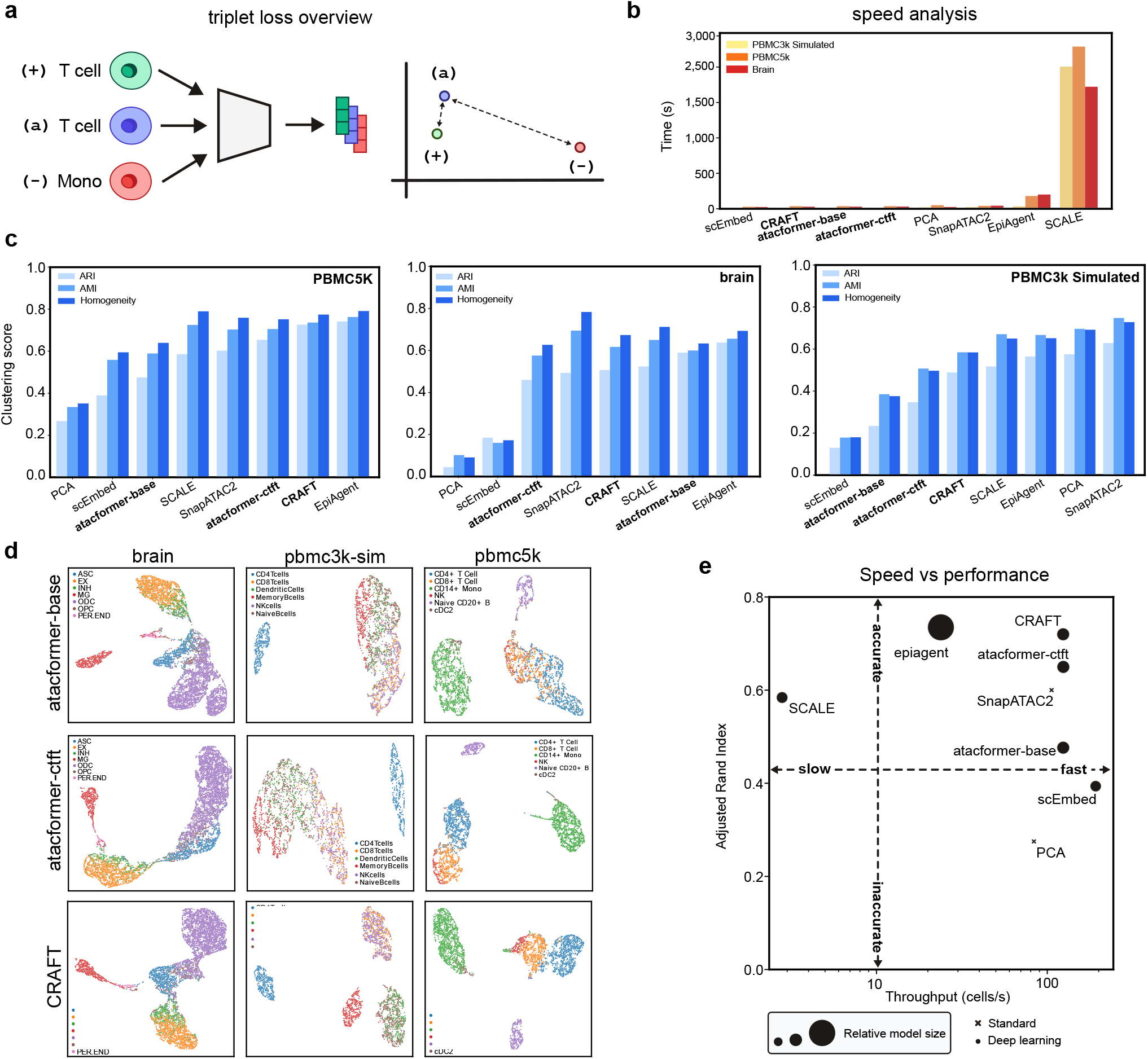
Atacformer clusters new scATAC data accurately in a zero-shot approach. **a**. Schematic of the supervised cell-type fine-tuning task using triplet loss. **b**. Runtime comparisons between Atacformer and other popular methods for clustering scATAC-seq data. **c**. Clustering performance of Atacformer-base and Atacformer-ctft on three separate PBMC datasets compared with other popular methods for clustering scATAC-seq data. **d**. UMAP plots highlight the latent space change when fine-tuning Atacformer on cell types. **e**. Runtime versus ARI chart for Atacformer and other popular methods.

To assess clustering efficiency and performance, we collected three separate datasets for evaluation: 1) A third-party, pre-annotated brain dataset; 2) a simulated PBMC dataset from bulk ATAC-seq data; and 3) a pre-annotated PBMC dataset. We generated cell embeddings for all cells in each dataset, and then used them to benchmark `atacformer-base, atacformer-ctft` and CRAFT against Principal Component Analysis (PCA) and several popular methods for scATAC-seq clustering^14,15,18,35^. To establish Atacformer’s capabilities for clustering data it’s never seen before (i.e. zero-shot clustering), all datasets were explicitly excluded from the training data for all three models.

Atacformer was one of the fastest methods evaluated; second only to our previous Word2Vec-based method scEmbed^35^ (Fig. 3B). The slowest methods were those that required the use of a very large model (EpiAgent, 1.5B parameters^18^), or required training from scratch on a combined dataset with both the original and new data (SCALE).

To evaluate clustering accuracy, we clustered the embeddings obtained from each method to obtain cluster labels. These labels were then compared to the ground-truth cell-type labels using three metrics: Adjusted Rand Index (ARI), Adjusted Mutual Information (AMI), and Homogeneity score (see Methods). As expected, the `atacformer-base` model underperformed other methods for PBMC-based datasets. The cell-type fine-tuned model increased clustering performance according to ARI by *≈* 15% across the PBMC datasets (Fig. 3C). These gains were also reflected in the training dataset for the fine-tuning procedure (Fig. S4). Notably, `atacformer-base` performed exceptionally well on the brain dataset, surpassing the cell-type fine-tuned model. We reasoned that this was because the cell-type fine-tuned Atacformer model was fine-tuned *only* on blood data, hurting its performance. Finally, we found CRAFT to perform very well across the datasets, highlighting the power of contrastive learning for embedding quality improvement. To further explore how the fine-tuning step improved the base model for this task, we compared the embeddings of the two Atacformer models visually using UMAP. The clustering score gains are clearly reflected in the UMAPs. For example, the cell-type fine-tuning approach significantly improves Atacformers ability to cluster the NK-cells along with differentiating between CD4+ and CD8+ T cells (Fig. 3D).

In practice, it is common for individual cells to yield more fragments than the model’s maximum context window, resulting in more tokens than can be processed in a single forward pass (Fig. S5A). We addressed this with two strategies: (1) truncating after the first C tokens (Fig. S5B), and (2) randomly sampling *C* tokens per cell (Fig. S5C). We adopted random sampling, reasoning that it better preserves the underlying biological heterogeneity and avoids systematic positional bias. To quantify the effect of context window truncation, we systematically varied the number of tokens sampled per cell and assessed the impact on clustering performance. Remarkably, we observed that Atacformer maintains strong clustering accuracy even with a substantial reduction in context window size (Fig. S5D). The performance remains robust until severe truncation, after which clustering metrics rapidly decline (Fig. S5E). This result suggests that the model is resilient to moderate input reduction, enabling efficient analysis without significant loss in biological resolution.

We highlight CRAFT’s striking performance when considering both speed and clustering ability together. For the PBMC5k dataset, CRAFT not only exhibited the best clustering performance, it also generated those clusters most quickly (Fig. 3E), yielding the highest cells-per-second throughput of all methods. In addition, we performed a preliminary batch-correction analysis across multiple PBMC datasets. Visually, Atacformer embeddings integrated as well as EpiAgent’s (Fig. S6), despite the latter’s much larger parameter count. This suggests that Atacformer’s zero-shot capabilities hold promise for rapid, multi-dataset integration without retraining, a key requirement for scalable single-cell analysis.

Finally, we omitted ChromFound^17^, a newly published scATAC-seq foundation model, from these panels since the code is not opensource and ARI/runtime on these exact PBMC benchmarks aren’t reported. In their work, they report ARI=0.48 on a similar PBMC dataset, while maintaining a throughput of 4 cells per second. This would make it the second slowest method benchmarked.

### Atacformer learns global regulatory structure in bulk region set data

Single-cell assays are redefining chromatin biology, yet aggregate profiles from bulk ATAC-seq, ChIP-seq, and DNase-seq are still far more common, and will likely continue to be produced in the future due to dramatically lower cost, as they provide useful information for homogeneous samples or when cell-to-cell heterogeneity is not important. In recent years the number of BED files uploaded to the Gene Expression Omnibus have increased by more than 10,000 files per year, most of them from bulk experiments of many varieties^36,37^. To help manage and interpret this growing archive, we recently developed BEDbase – a unified platform for aggregating, analyzing, and serving genomic region data. We therefore asked whether Atacformer, trained on single-cell data, could generalize to the more heterogenous and voluminous profiles posted to GEO, and thereby provide practical utility for BEDbase, such as by imputing missing key metadata or enhancing region annotations.

To assess this, we fine-tuned Atacformer on BED files annotated with hg38 on BEDbase. We downloaded and tokenized over 35,000 BED files and continued pretraining `atacformer-base` using ELECTRA-style token replacement prediction for 10 epochs, yielding `atacformer-bb`. Using `atacformer-bb`, we generated embeddings for BED files from the top 10 cell lines and assays in our bulk data training set. The embeddings clustered by assay type (Fig. 4A), suggesting that the model captured assay-specific signal features, as well as by cell line (Fig. 4B), with individual clusters corresponding to cell lines, indicating the model learned general patterns of regulatory biology.

**Figure 4.**
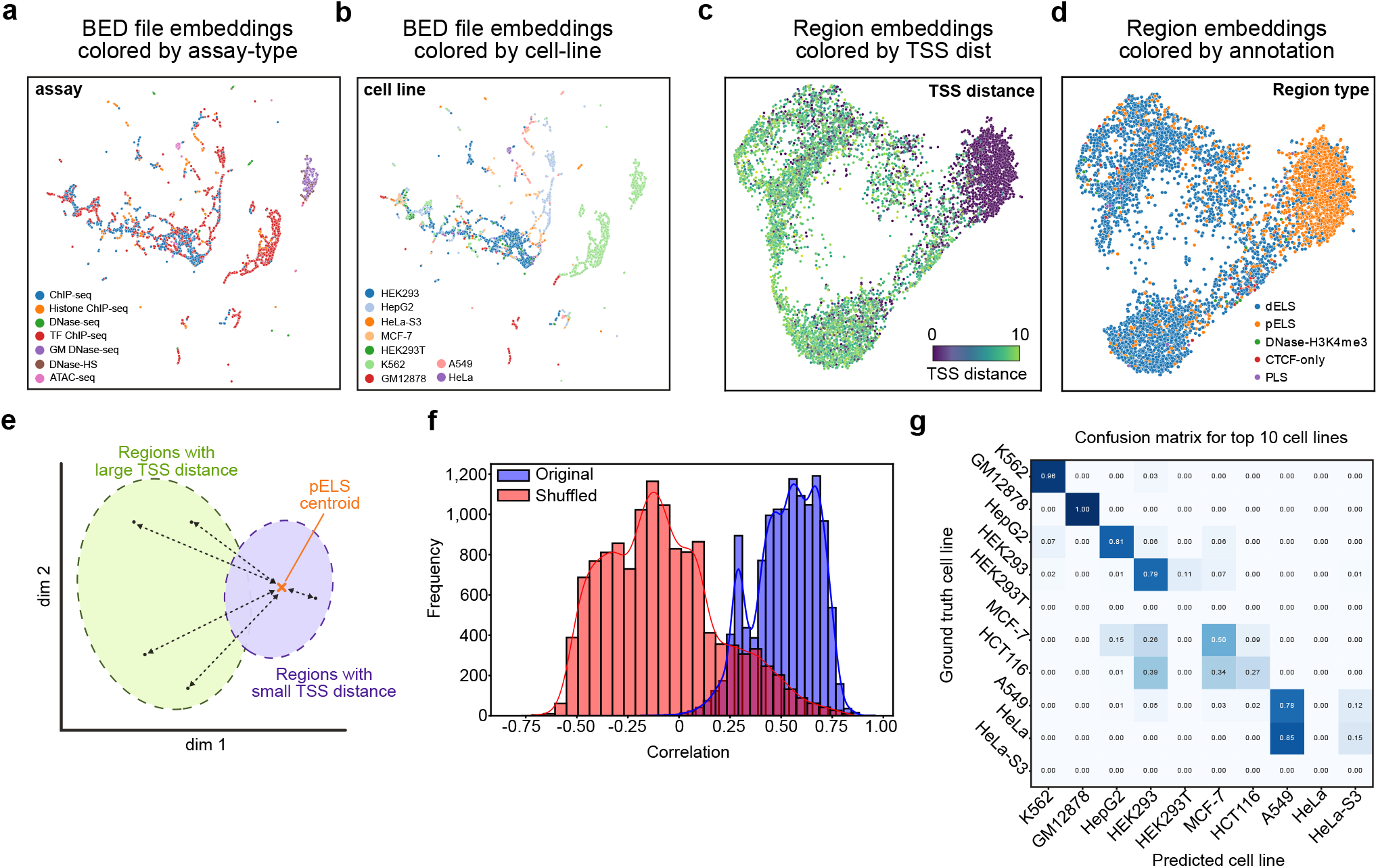
Atacformer generalizes to bulk regulatory datasets. **a**. UMAP projection of ≈ 10,000 BED file embeddings from BED files on BEDbase. Points are coloured by assay, revealing discrete clusters of assay types. **b**. The same UMAP projection colored by cell line shows distinct groupings for common cell types, indicating that Atacformer encodes both experimental and biological context without supervision. **c**. UMAP visualization of a set of contextualized region embeddings from an example BED file, colored by TSS distance. **d**. The same UMAP visualization, instead with each region embedding colored by its annotation from the ENCODE SCREEN project dELS = distal enhancer-Like; pELS = proximal enhancer-like; DNase-H3k4Me3 = high H3K4me3, low H3K27ac; PLS = promoter-like. **e**. Schematic of the global TSS distance analysis. The distance of each contextualized region embedding to the pELS centroid is found. In general, regions with a high TSS distance will have a larger distance. **f**. Distribution of Pearson correlation coefficients when correlating a region’s annotated TSS-distance to its distance to the pELS centroid. **g**. Confusion matrix for our cell line classifier, limited to the top ten cell lines in the bulk dataset.

To investigate whether token-level embeddings also reflect learned biology, we annotated each of the model’s 890,704 regions with its distance to the closest transcription start site (TSS) and its detailed region classification according to the ENCODE registry of cCREs (see Methods). We reasoned that the contextualized region embeddings produced by the transformer would enable rich information to be encoded in the embeddings. When visualizing an example BED file with UMAP, we found that contextualized embeddings clustered by TSS distance (Fig. 4C). To confirm that the model is distinguishing promoters from enhancers, we also colored the contextualized region embeddings by their region annotations, and found that regions clustered broadly into groups of proximal enhancer-like sequences (pELS) and distal enhancer-like sequences (dELS; Fig. 4D).

We next sought to interrogate whether this trend held broadly for all BED files in the training dataset. For this, we developed a novel assessment score inspired by our previous work on region embedding evaluation^38^ called the TSS distribution score. Briefly, the score is computed for each region using three steps. First, we identify the pELS centroid in the embedding space. Second, each region’s embedding distance to this centroid is computed. Finally, we correlate each region’s TSS-distance to its distance to the pELS centroid. We hypothesized that regions close to the centroid would also have corresponding small TSS distances, whereas regions far away from the centroid would have corresponding large TSS distances (Fig. 4E). As a control, we also computed the TSS distribution score for shuffled labels. We found that the contextualized embeddings for BED files consistently had TSS distribution scores greater than zero, while controls yielded a distribution of scores centered at zero (Fig. 4F). This result confirms that the contextualized embeddings are capturing learned regulatory biology across cell lines and assay types.

Finally, we sought to use Atacformer to help annotate data on BEDbase (Fig. S7). Many files in the database are missing critical metadata – particularly cell line annotations. We therefore hypothesized that Atacformer could be used to impute this missing data and built a proof of concept pipeline to annotate missing cell line information for all BED files. Specifically, we hypothesized that the BED-file embeddings could be used as input to a classifier to predict the missing annotation.

To build the pipeline, we split our dataset into two groups: 1) BED-files with annotated cell lines and 2) BED-files with cell line annotations missing, but inferrable from the file description (see Methods; Fig. S8A,B). We use the BED-files with annotated cell lines to train an XGBoost decision tree model^39^, and then subsequently use the trained model to predict the cell lines for the BED-files with missing annotations, using the inferred annotations as ground truth labels for evaluation. The decision tree model was effective at annotating the missing cell lines. The model achieved an F1 score of 0.85 and an accuracy of 86% across over 275 cell lines (Fig. 4G). Taken together, these results highlight how Atacformer is able to generate powerful embedding representations of bulk genomic interval data with practical downstream use.

### Direct raw-fragment processing with Atacformer accelerates scATAC analysis

Existing scATAC-seq deep learning models such as EpiAgent, ChromFound, and SCALE require count matrices as input. This necessitates a pre-processing step using a tool like ArchR or SnapATAC2^12,14^ to generate the necessary input format when working with fragments files. In contrast, because of its tokenization procedure, Atacformer is uniquely positioned to generate single-cell embeddings immediately from fragments files. By leveraging interval-overlap-based tokenization, Atacformer bypasses the need for tedious preprocessing steps and produces embeddings directly from fragments, enabling more flexible pipelines and zero-shot analyses. This can improve both speed and flexibility in analysis pipelines (Fig. 5A).

**Figure 5.**
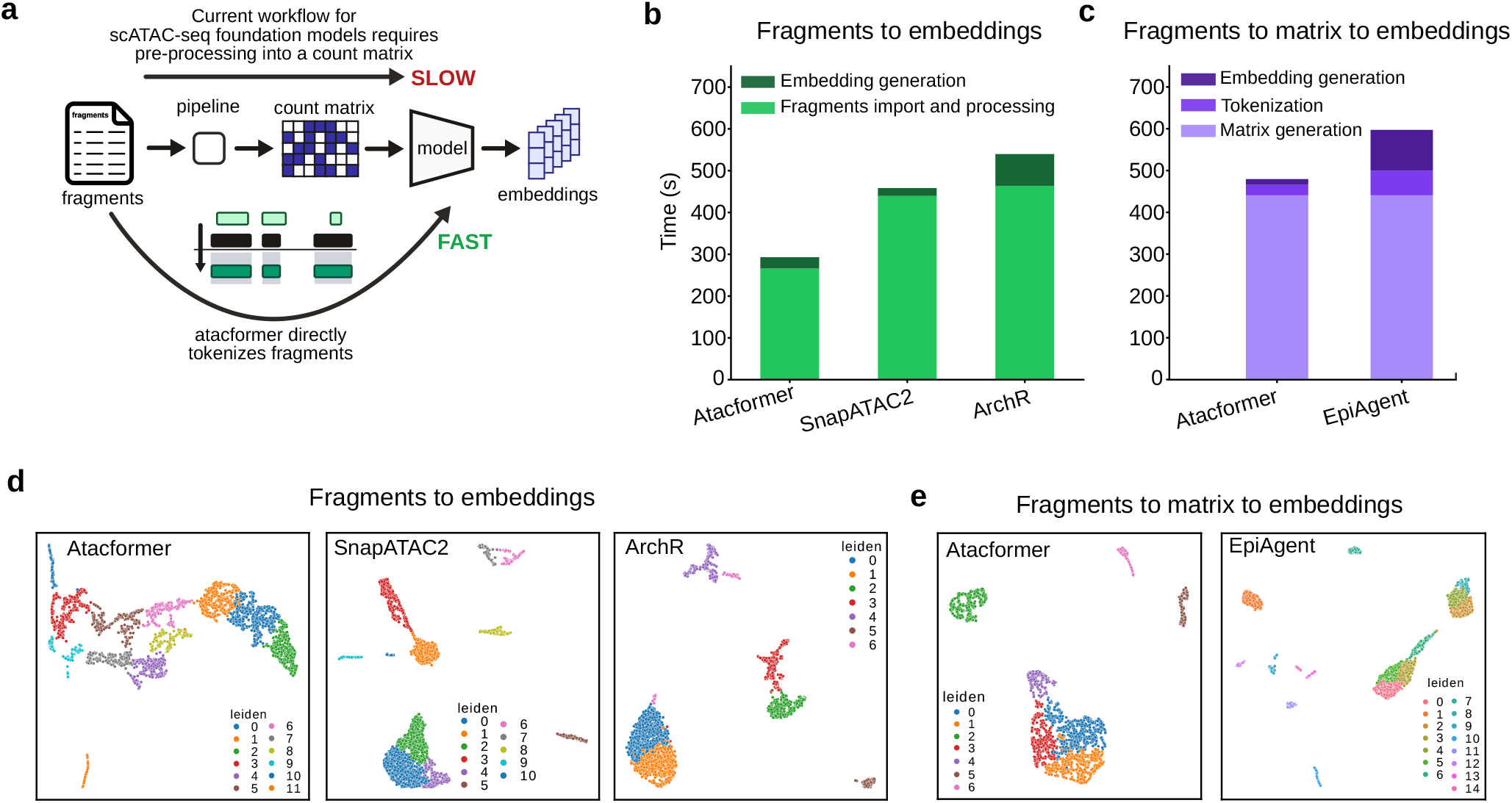
Atacformer is the only method that operates on sequence fragments directly. **a**. Schematic showing the difference between Atacformers ability to generate embeddings for fragments files and other current scATAC-seq foundation models. **b**. Speed comparison of processing times for Atacformer with SnapATAC2 and ArchR when processing direct fragments files. Fragments file processing is grouped into two main steps: 1) fragments file importing and 2) cell embedding generation. **c**. Speed comparisons between Atacformer and EpiAgent count matrix processing. Count matrix processing is grouped into three main steps: 1) matrix generation, 2) tokenization, and 3) embedding generation. **d**. UMAP visualizations for single-cell embeddings starting with fragments files for ArchR, Atacformer, and SnapATAC2. **e**. UMAP vsualizations of single-cell embeddings starting with count matrices.

To examine this, we obtained a new 3,000 cell dataset from flash-frozen human healthy brain tissue from 10X genomics. We call this *brain3k*. We then conducted two separate analyses: 1) fragments to cell embeddings, and 2) count matrices to cell embeddings. For the first analysis, starting from the fragments file, we processed the data in three ways: 1) using Atacformer and its native tokenizers; 2) using SnapATAC2; and 3) using ArchR. We organized this processing into two main steps: 1) fragments import and filtering, and 2) embedding generation. We measured the time it took to conduct each step for each method (see Methods). We then used the resultant embeddings to generate UMAP visualizations. For the second analysis, we took the processed count matrices and fed them to Atacformer and EpiAgent, generating embeddings and corresponding UMAP visualizations, similarly measuring the time it took to conduct each step.

We first used the Atacformer tokenizers to import and tokenize the fragments, filtering by barcodes. We then used these tokenized fragments to generate cell-level embeddings for each cell. In timed comparisons, the end-to-end wall time for Atacformer – comprising fragments import/filtering and embedding generation – was substantially lower than either SnapATAC2 or ArchR (Fig. 5B). Most of the runtime in the baseline pipelines was spent on I/O-heavy import and matrix construction, whereas Atacformer’s Rust-backed interval-overlap tokenization minimizes this overhead by operating directly on the fragments file^28^.

Then we next sought to use the processed count matrices as input into Atacformer and compare the performance against EpiAgent, a recently published scATAC-seq foundation model. We found that the majority of the processing time for both methods was spent on pre-processing the count matrix into a suitable input format. However, once this step was complete, Atacformer generated embeddings substantially faster than EpiAgent. In particular, the Atacformer tokenization and embedding generation steps were approximately 2.5x faster than EpiAgent’s equivalent steps (Fig. 5C).

Using the resulting embeddings from the fragments and count matrices, we visualized cells with UMAP and performed Leiden clustering. For the fragments-based analysis, Atacformer produced coherent, well-separated clusters that were comparable to or better resolved than those obtained from matrices built with SnapATAC2 or ArchR (Fig. 5D). Notably, Atacformer achieves this in a zero-shot manner – without peak calling or an intermediate peak-by-cell matrix - by leveraging fragment co-occurrence structure within cells. For the count-matrix-based analysis, Atacformer again produced well-separated clusters that were comparable to or better resolved than those from EpiAgent (Fig. 5E).

Together, these results demonstrate that Atacformer enables practical, faster, and more flexible scATAC-seq workflows: embeddings can be generated directly from fragments, eliminating prerequisite matrix construction and supporting downstream analyses (e.g., clustering) with minimal preprocessing.

### Contextualized region embeddings from scATAC-seq data infers cryptic TSSs

A strength of the Atacformer architecture is its ability to generate token-level embeddings for individual cis-regulatory elements (cCREs). Unlike other foundation models that produce cell-level representations, Atacformer enables direct investigation of relationships between discrete, well-annotated, genomic regions within a single cell. Building on our initial exploration with bulk ATAC-seq data, we applied this approach to single-cell data, capturing contextualized embeddings for each region prior to aggregation into cell-level representations (Fig. 6A).

**Figure 6.**
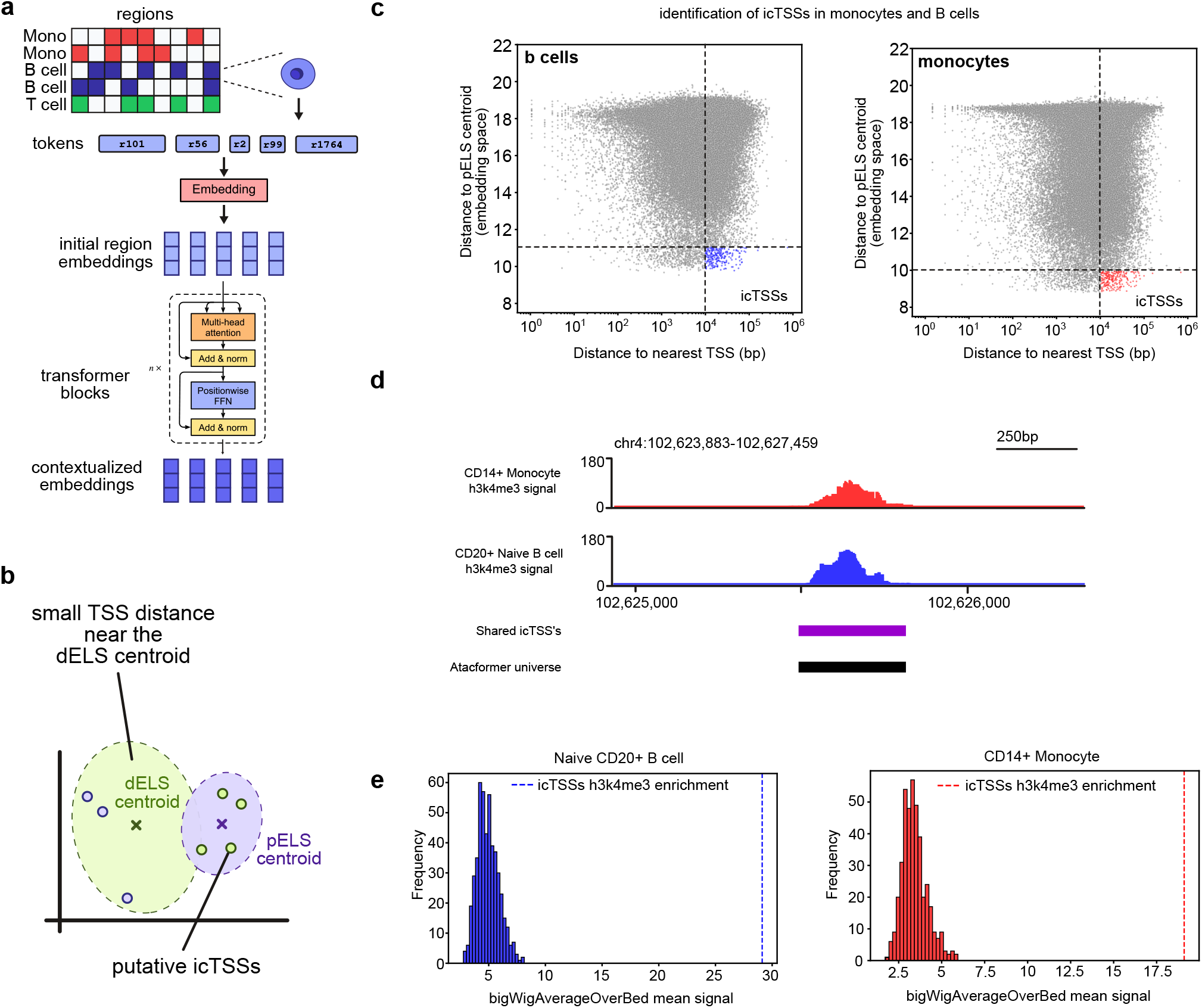
Atacformer uncovers weak promoters using scATAC-seq data alone. **a**. Schematic of generating contextualized region embeddings from a single cell. **b**. Schematic of the canonical contextualized latent space for co-accessible regions in a single-cell. In general, distal enhancer-like sequences cluster together while proximal enhancer-like sequences cluster together. **c**. Plot of annotated TSS distance versus pELS centroid distance. We highlight paradoxical regions which are both annotated as very far from the TSS and found very close to the pELS centroid. **d**. Example of an icTSS with strong H3K4me3 signal despite no annotated promoter nearby. **e**. Histograms of H3K4me3 enrichment in a randomly generated background distribution compared to the icTSSs in both CD14+ monocytes and CD20+ Naive B cells.

As with bulk data, single-cell region embeddings were broadly structured by annotation, clustering according to TSS distance and their classification as proximal (pELS) or distal (dELS) enhancer-like (Fig. 6B). Within this structure, however, we identified a discordant subset of regions. These elements were annotated as distal from any TSS, yet their embeddings clustered with the pELS centroid – a location dominated by promoter-proximal regions. We hypothesized that this discrepancy reflects functionally important sites, such as weak promoter regions, which we term inferred cryptic TSSs (icTSSs; pronounced “iced-teas”). We use ‘cryptic’ to denote TSSs hidden from current genome annotations, which may include both epigenetically repressed sites and active but unannotated promoters^40^.

To investigate this, we subsampled 10,000 single-cells from the Luecken2021 dataset for CD14+ monocytes and naïve CD20+ B cells. For each cell, contextualized co-accessible region embeddings were generated, and each region’s distance to the nearest TSS was plotted against its distance to the pELS centroid. Both cell types exhibited a modest but positive correlation between these metrics (Spearman’s *ρ* = 0.30 and 0.29, respectively). We then defined icTSSs as regions annotated as highly distal from a TSS yet paradoxically embedded very close to the pELS centroid (Fig. 6C).

To validate that icTSSs represent latent promoter regions, we downloaded ChIP-seq data for H3K4me3, a canonical promoter marker, for both cell types^41^. We reasoned that H3K4me3 signal suggests a region harbors promoter activity. We found that icTSSs were strongly enriched for H3K4me3 (Fig. 6D; Fig. S9). To assess statistical significance, we constructed null distributions by repeatedly sampling random region sets from the Atacformer vocabulary and comparing their signal overlap with H3K4me3 (see Methods). Against this background, monocyte-specific icTSSs showed a 6.33-fold enrichment for monocyte H3K4me3 peaks, while B-cell icTSSs showed 6-fold enrichment for B-cell peaks (empirical p < 0.001 in both cases) (Fig. 6E). Together, these results demonstrate that Atacformer’s contextualized embeddings can identify bona fide weak promoters directly from ATAC-seq data, and that this signal is cell-type specific.

## Discussion

Atacformer introduces a general-purpose transformer-based foundation model for chromatin accessibility data, demonstrating strong performance across a diverse set of tasks, including cell-type clustering, fragment file processing, bulk ATAC-seq embedding, and multimodal integration with RNA-seq. Unlike prior models, Atacformer explicitly tokenizes genomic intervals as discrete units and discards reliance on positional encodings, instead encouraging the model to learn contextual biological relationships directly from data. This aspect of Atacformer directly builds on our previous work with learning genomic region embeddings^42,43^.

One major advantage of Atacformer’s region-based approach is its ability to operate directly on raw fragment files, bypassing the need for intermediate matrix generation. This reduces processing time, making it especially valuable for large-scale or time-sensitive analyses. Compared to existing scATAC pipelines, Atacformer achieves comparable biological fidelity while accelerating analysis by 80% in our benchmarks. This positions Atacformer as a fundamentally streamlined alternative to conventional workflows.

Our results show that Atacformer’s pre-training strategy, based on ELECTRA-style replaced token detection, enables generalization across both single-cell and bulk data. Fine-tuning on BEDbase bulk datasets reveals that Atacformer embeddings encode biological information such as assay type and cell line identity even in the absence of labels. Token-level embeddings are organized by promoter-enhancer distance without ever having seen explicit TSS annotations during training, underscoring the model’s ability to infer global regulatory structure from chromatin accessibility alone.

By integrating Atacformer with Geneformer, we further demonstrate Atacformer’s extensibility to multimodal contexts. The CRAFT framework highlights that chromatin-accessibility embeddings can align with transcriptomic signals in a shared latent space, enabling cross-modal retrieval and RNA imputation from ATAC alone. This not only expands Atacformer’s applicability to joint profiling datasets, but also opens the door for future extensions such as natural language and epigenome integration or DNA methylation-RNA alignment using similar dual-encoder strategies.

While Atacformer achieves strong results with fewer parameters than other models, several limitations remain. First, the lack of positional encoding–while intentional–may hinder tasks requiring spatial resolution, such as enhancer-promoter linking across large genomic distances. Second, our approach to fragment tokenization is sensitive to the predefined vocabulary and resolution, which could affect generalization across genome builds or non-human species. Finally, our evaluations, particularly in multimodal alignment, were limited to well-annotated datasets; broader assessments in low-quality or noisy settings are needed.

Looking ahead, several avenues for extending Atacformer’s capabilities are promising. These include: 1) Longer context windows or streaming architectures for encoding ultra-complex single-cell profiles; 2) Generative pre-training approaches, such as diffusion or masked span prediction to enable more flexible inference tasks; and 3) Application to clinical and diagnostic datasets, especially in cancer, where chromatin structure is often perturbed.

In conclusion, Atacformer serves as both a performant model and a software framework for ATAC-seq analysis. Its general-purpose embeddings, fragment-level input pipeline, and compatibility with other models make it a powerful tool for epigenomic research. Importantly, Atacformer’s contextualized region embeddings reveal functionally active but unannotated promoter elements, demonstrating how foundation models can uncover hidden biological features directly from chromatin accessibility data. By bridging bulk and single-cell assays, integrating modalities, and enabling fast and interpretable analysis, Atacformer contributes to the growing ecosystem of foundation models in biology and offers a blueprint for future advances in genomic machine learning that not only improve computational performance but also deepen our understanding of gene regulatory mechanisms.

## Methods

### Data collection and pre-processing

To develop a large training dataset of single-cell ATAC-seq data, we identified, downloaded, and uniformly processed data from three main sources: 1) the Gene Expression Omnibus, 2) the Human Cell Atlas, and 3) the 10X genomics dataset repository. Detailed information on dataset contents and availability can be found in supplemental. All datasets, unless noted, were downloaded as raw `.fastq` files. We designed and built a multi-stage pipeline to uniformly process the `fastq` files. First, we utilized CellRanger ATAC 2.1.0 to convert the fastq files into processed fragment files. Next, each fragment file was imported and initially processed using SnapATAC2^14^. We utilized all recommended parameters noted in the “atlas-scale analysis” tutorial (“https://kzhang.org/SnapATAC2/tutorials/atlas.html“). Both steps were parallelized on our computing cluster using the `looper` and `PEP` framework^44^.

Next, we again leveraged SnapATAC2 to perform atlas-wide dimensionality reduction, batch correction, and clustering. We performed a two-stage clustering approach. First, a coarse clustering across the entire dataset using Leiden clustering^45^ (Fig. S10), and then a secondary intra-cluster clustering also using leiden within each cluster (Fig. S11). This yielded 359 distinct single-cell clusters within the atlas across all datasets. Each of the 359 clusters was pseudo-bulked into separate `.fragment.tsv` files for downstream analysis.

### Generation of a uniform model vocabulary

A uniform vocabulary is essential for genomic region tokenization. For this, we leveraged both public and previously developed tools by our lab for generating genomic interval consensus sets. We utilized `macs3` to perform peak-calling^46^ on each of the 359 pseudo-bulked fragment files, resulting in 359 corresponding `.narrowPeak` files. Specifically, we used the peakcall function with the following parameters: `-g hs -f BEDPE -q 0.01 --nomodel --shift -75 --extsize 200 --keep-dup all –B --SPMR`.

We next utilized our tool `uniwig` to unify all 359 peak sets into big wig (`.bw`) coverage tracks for the start, cores, and ends of all called peaks across all clusters (citation). For `uniwig` we used the following parameters: `-m 5 -s 1 -y wig -z 2`. This resulted in three coverage track files for the starts, cores, and ends.

Finally, we used the coverage tracks as input into our previously published universe creation methods in `geniml`^27^. Using both the coverage cutoff and hidden markov model (HMM) algorithms, two consensus sets were created. These two bedfiles were finally merged into one unifying vocabulary for the model using `bedtools merge`^47^.

The final vocabulary has 890,704 distinct genomic regions into which all cells are tokenized.

### Genomic interval tokenization

We conceptualized a unique tokenization method for our models that is designed to be as flexible and simple as possible, enabling broad use of Atacformer for many data types including bulk-ATAC seq data. Any entity that can be represented as a BED-file is valid input to the model. As an example, a single-cell from a scATAC-seq experiment can be thought of as a “bag of co-accessible regions”consisting of a few thousand open chromatin regions. Each of these regions is overlapped with the model’s pre-defined vocabulary and serves as input to the embedding module and subsequent transformer encoder.

We leverage two highly-efficient methods for interval overlap computation: AIList^48^ and BITS^49^. We’ve implemented both algorithms in Rust and have made them available in Python for in-memory tokenization with a huggingface-compatible API. Code and documentation for our tokenizers can be found on GitHub in our gtars crate/package “https://github.com/databio/gtars“.

### ELECTRA pre-training

To pretrain Atacformer on single-cell data, we employed an ELECTRA-style replaced-token detection strategy. While most transformer encoder models use masked language modeling (MLM) for self-supervised pretraining^50^, we found that MLM was poorly aligned with the properties of our model and data modality in two key ways. First, MLM requires computing a full probability distribution over the vocabulary at each training step. With nearly 1 million tokens, this becomes computationally intractable. Only recently have methods emerged to address this issue. Motivated by the growing vocabulary sizes in modern large language models (LLMs), techniques like Liger kernels^51^ and Cut Cross Entropy^52^ use linear-time approximations of cross entropy to dramatically reduce space and time complexity. We found these strategies applicable to Atacformer as well, since MLM is fundamentally a token prediction task. However, a second, more fundamental problem emerges: we identified that MLM effectively enforces an order among masked tokens, while shuffling the tokens should have no effect on the information content of a single-cell or corresponding regionset. More specifically, we recognized that the model predicting the correct tokens, but *out of order* was common and would provide an incorrect training signal to the model.

ELECTRA side-steps both of these problems entirely by reframing the pre-training task as binary classification: for each token, the model predicts whether it was replaced or not. This approach does not depend on the model predicting tokens in a specific order, and moreover, doesn’t require computing a probability for all 890K tokens. Although genomic coordinates offer a natural means to introduce sequence order, we sought to avoid the model overly relying on deterministic positional cues, instead incentivizing it to capture meaningful biological patterns and relationships. To that end, Atacformer is distinctly free of any form of positional information in its input embeddings.

### Formal specification of tokenization and pre-training

We begin by fixing a **global vocabulary** *U* of non-overlapping genomic regions derived from our consensus universe (see previous section):

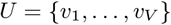

For each single cell *c* we observe an unordered set of raw, **co-accessible** regions

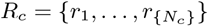

Tokenization reduces these raw intervals to their canonical vocabulary representatives via a simple interval intersection:

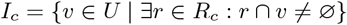

We then map each vocabulary element to its integer identifier, producing a sequence of token indices

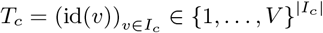

To create a learning signal we apply **ELECTRA-style corruption**. Each position *j* is independently selected for replacement with probability *p* = 0.45:

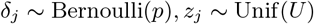

The corrupted token sequence is therefore:

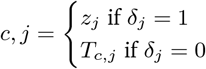

Every token id is looked up in a shared embedding matrix *E* ∈ ℝ^*V* × *d*^ to obtain dense vectors

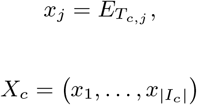

These embeddings pass through *L* stacked transformer encoder layers (no positional encodings are supplied):

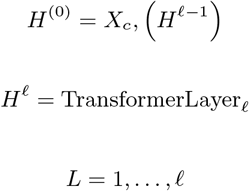

The final contextual embedding at each position *j* is

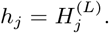

A lightweight classifier head converts each contextual vector into the probability that the original token was **replaced**:

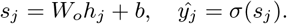

Training minimizes the binary cross-entropy **replaced-token detection** loss over all positions in the cell:

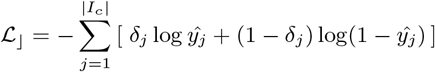

Averaging ℒ _⌋_ over the mini-batch and optimizing with AdamW completes the pre-training procedure.

### Cell embedding calculation

To generate single-cell embeddings, we first tokenize a single-cell according to the steps outlined above. We then pass the tokens through the model to obtain a contextualized embedding representation for each region-token in that cell:

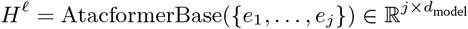

Where *e*_*j*_ is the initial token embedding for the *j*^th^ region token. We then obtain cell-embeddings by pooling all contextualized region-embeddings through mean-pooling:

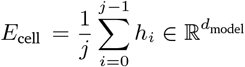

where *h*_*i*_ is the contextualized embedding vector for the *i*^th^ token.

### Triplet loss calculation

For cell-type fine-tuning we use a standard triplet-loss formula. For each training step, the model sees three cells: 1) an anchor cell which may be of any cell-type, 2) a positive example which is of the **same** cell-type as the anchor, and 3) a negative example which is of **another** cell-type than the anchor cell. We pass each cell through the model and mean-pool token embeddings to obtain a single embedding to represent the entire cell; one for the anchor (*a*), the positive (*p*), and the negative (*n*). Loss for a single mini-batch is computed as such:

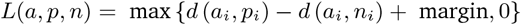

where

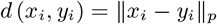

We use the torch module `torch.nn.TripletMarginLoss` with default values, margin = 1.0, and *p* = 2.0.

### Datasets for clustering evaluation

#### PBMC5k NextGEM v 1.1

For PBMC5K, we obtained raw matrix, peak, and barcode files from the 10X website: “https://www.10xgenomics.com/datasets/5-k-peripheral-blood-mononuclear-cells-pbm-cs-from-a-healthy-donor-next-gem-v-1-1-1-1-standard-2-0-0”. These three files were converted to an `AnnData` object from the `scanpy` package.

#### Brain dataset

For the pre-annotated brain dataset, we utilize a multi-omic single-nucleus study of 191,890 nuclei in late-stage Alzheimer’s disease (AD)^53^. Cells in this dataset were annotated using gene expression data to assign ground-truth labels to each cell. These labels were used for downstream clustering metrics evaluation.

#### Simulated

Evaluating model performance on real, pre-annotated datasets is subjected to the bias in the annotation procedure. This can cause misleading results according to inaccuracies in the labeling process. To that end, we supplemented our two datasets with a third, simulated dataset. Following a similar procedure to Chen *et. al*.^54^, we generated a simulated scATAC-seq dataset using bulk-ATAC data from ENCODE. We first generate a peak by count matrix from 5 bulk-ATAC seq datasets: NK Cells (ENCSR305QTE), Memory B Cells (ENCSR610AQP), Naive B Cells (ENCSR685OFR), Dendritic Cells (ENCSR237VSF), CD4+ T cells (ENCSR841LHT), and CD8+ T cells (ENCSR476VJY). We leverage the simulation code provided by the Pinello lab: https://github.com/pinellolab/scATAC-benchmarking/blob/master/Synthetic_Data/Simulate_scATAC/BoneMarrow/simulate_bonemarrow_depth.ipynb

#### PBMC dataset cell-type annotation

Because ground-truth labels are necessary for adequately assessing the clustering performance of cell-embeddings, we performed cell-type annotation on all three datasets. Each annotation was performed in an identical manner. To do so, we followed a very similar approach to the cell-type annotation approach used in scEmbed^35^. Briefly, we leveraged a pre-trained scEmbed model trained specifically on a high-quality blood dataset, Luecken2021^32^. Embeddings were generated for both the reference dataset (Luecken2021) and the query datasets (PBMC 1/5/10k). Then, using the shared latent space, we performed a K-nearest-neighbors (KNN) label transfer task. We used `scEmbed` from the `geniml` module on GitHub: “https://github.com/databio/geniml” and the `KNeighborsClassifier` from `sklearn.neighbors`. Due to the intrinsic sparsity of many detailed T-cell subtypes, we collapsed these rare variants into broader T-cell categories. This aggregation prevents overfragmentation during clustering, ensuring a more statistically robust and biologically meaningful representation of T-cell populations.

#### Clustering algorithm

We leveraged K-Means clustering for all evaluation tasks. We use the scikit-learn implementations. Because ground truth labels were known, we used the number of unique cell-types to set the number of clusters to generate. All other parameters were set to their respective defaults.

### Clustering metrics

#### Adjusted rand index

The ARI is a metric for evaluating the similarity between two data clusterings. This is achieved by counting pairs that are assigned to the same cluster label. Mathematically, it is computed by:

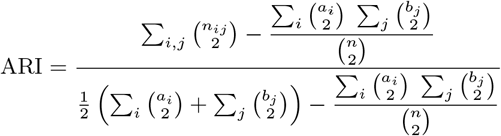

where *n*_*ij*_, *a*_*i*_, *b*_*j*_ are diagonal values, row sums, and column sums respectively from the contingency table that describes the frequency distribution of the cluster labels from ground-truth and predicted clusterings. We use the adjusted rand score function from the `scikit-learn` python package.

#### Adjusted mutual information

The AMI, intuitively, is a measure of the amount of information that two clusterings share. It’s used to evaluate how well two clusterings agree with each other^55^. We compute AMI through the scikit-learn package using the adjusted mutual info score function.

#### Homogeneity score

The homogeneity score is an entropy-based external cluster evaluation metric that measures how far from perfect an incorrect clustering solution is^56^. We employ the scikit-learn homogeneity score function to measure this metric for each dataset.

#### UMAP visualization

UMAP projection provides a quick way to visualize global structure in the embeddings. For all UMAP visualizations, we utilize the `umap-learn` package in python and use default parameters.

### Fragment file ingestion and tokenization

Fragment files are ingested using our gtars package.

### Fragment agreement evaluation metrics

#### Processing time

To measure processing time, we leverage the python standard library `time` module.

### Bulk training data selection

To generate a large dataset for fine-tuning on bulk data, we first started with all bed-files annotated with hg38 on BEDbase. Because Atacformer has a context window of 8,192 tokens, we next filtered down these bedfiles into a subset that could reasonably fit within this context window, subsampling tokens as necessary. We set the cutoff for the number of regions in the bedfile to be 81,920 (10x the context window).

We tokenized the BED files that met this criteria and used them as input into the training pipeline, subsampling tokens from the file when necessary.

### TSS distance annotation of our universe

To annotate the distance to the nearest TSS to each token in our vocabulary, we first downloaded the most recent comprehensive gene annotation (GTF) file from GENCODE (“https://www.gencodegenes.org/human/release_38.html”). We filtered this file to obtain just the TSS annotations using common unix command-line tools like `awk` and `sort`.

Next we leveraged `bedtools` to obtain distances to the nearest TSS. Specifically, we used the `bedbase closest` command with the `-t first` flag to ensure each region in our universe was only associated with one TSS.

### Spearman correlation

The spearman correlation can be computed as follows:

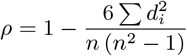

where *d*_*i*_ is the difference between the two ranks of each observation and *n* is the number of observations. To compute the value, we leveraged the `scipy.stats` module and the `spearmanr` function.

### Multiome data processing

To curate a large multiome dataset, we downloaded and processed four datasets: three from the 10X genomics dataset repository and then the previously described Luecken2021 dataset^32^. The three 10X datasets were: 1) brain3k multiome, 2) kidney22k multiome, and 3) pbmc10k multiome (Table S2). For each dataset, we downloaded the cell by feature matrix as a matrix-market file (`.mtx`), the barcodes as a `.txt` file, and the features as a `.tsv` file. We combined these files into an `.h5ad` file for each dataset using the `scanpy, pandas` and `scipy` python packages.

Each dataset was tokenized into the universe as previously described and used for training CRAFT.

### CRAFT architecture

The CRAFT architecture closely follows the design of the CLIP model^30^, which jointly trains two separate encoders to project different modalities into a shared latent space. In our implementation, we replaced the original CLIP encoders with domain-specific architectures: the ATAC encoder was substituted with the Atacformer, a transformer-based model tailored for chromatin accessibility data, and the RNA encoder was replaced with geneformer, optimized for gene expression profiles.

We train CRAFT starting with a pre-trained Geneformer and Atacformer model. Namely, we use `Geneformer/gf-12L-30M-i2048` and `databio/atacformer-base-hg38` respectively. We trained for 15 epochs using a linear learning rate scheduler with a maximum learning rate of 5e *−* 5.

### pbmc5k dataset processing for RNA-imputation experiments

To prepare a dataset, we closely follow the “RNA integration” tutorial offered by the SnapATAC2 “Annotating cell clusters” tutorial. Briefly, the pbmc5k dataset was imported from SnapATAC2, filtered, dimensionality-reduced, and subsequent cell-type annotation was performed using scvi.

### CRAFT RNA decoder

Using `pytorch`, we built a small decoder to predict a cell’s RNA-expression profile from its corresponding shared latent space ATAC embedding. We used a simple feedforward neural network with one hidden layer.

To obtain the shared latent space embedding from the ATAC data, we first encoded the cell’s ATAC profile using an encoder network. The encoder consisted of a fully connected layer that projected the high-dimensional ATAC input into a lower-dimensional latent space, followed by a non-linear activation function (ReLU). The output of this encoder served as the input to the RNA decoder.

The overall architecture thus consisted of an ATAC encoder, which mapped the input ATAC features to a latent representation, and an RNA decoder, which predicted the RNA expression profile from this latent space.

### Annotation of Atacformer universe for TSS distance and region type

To annotate the distance to the nearest TSS to each token in our vocabulary, we first downloaded the most recent comprehensive gene annotation (GTF) file from GENCODE release 38. We filtered this file to obtain just the TSS annotations using common unix command-line tools like `awk` and `sort.` Next we leveraged `bedtools` to obtain distances to the nearest TSS. Specifically, we used the `bedbase closest` command with the `-t first` flag to ensure each region in our universe was only associated with one TSS.

Similarly, we downloaded the latest cCRE annotations from ENCODE screen (https://screen.encodeproject.org) for hg38. We utilized `bedtools intersect` to annotate each region with a discrete label (pELS, dELS, CTCF, etc).

### H3K4me3 null distribution generation

To evaluate H3K4me3 signal enrichment in our icTSS regions across cell types, we first generated a null distribution representing the expected signal overlap in randomly selected genomic regions of comparable size. Specifically, we randomly sampled *N* regions from the Atacformer universe – where *N* equals the number of icTSS regions in each set (B cells and monocytes) – using standard Unix command-line utilities such as `shuf.` This sampling procedure was repeated 500 times to build a distribution of random region sets. For each set, we computed the mean coverage using `bigWigAverageOverBed` from the `bigtools` package^57^, then averaged the resulting signal across all regions. The resulting distribution was plotted as a histogram, and the same statistic was computed for the true icTSS regions to quantify their enrichment relative to the null.

## Supplemental material

**Supplementary Table S1.** Overview of datasets used in the pre-training of Atacformer.

| Dataset | Tissue | Disease state | No. cells | GSE/Link | Author |
| --- | --- | --- | --- | --- | --- |
| Human single-cell atlas | Atlas | Healthy | 615,998 | GSE184462 | Ren 2021 |
| Brain 107k | Brain | Healthy | 107,057 | GSE168408 | Lister 2023 |
| Atlas of tonsil | Tonsil | Healthy + Disease | 70,775 | - | Massoni-Badosa 2025 |
| Luecken2021 | Blood | Healthy | 69,249 | - | Luecken2021 |
| Parkinson's 65k | Brain | Disease | 65,589 | GSE148434 | Jung 2023 |
| Muscle 23k | Muscle | Disease | 23,593 | GSE174376 | Dyer 2022 |
| Kidney 22k | Kidney | Healthy | 22,772 | - | 10X |
| Cornea atlas | Eye | Healthy | 1,209 | GSE155683 | Lako 2021 |

**Supplementary Table S2.** Overview of multiome datasets used in the CRAFT fine-tuning process.

| Dataset | Tissue | Disease state | No. cells | Source |
| --- | --- | --- | --- | --- |
| Brain3k Multiome | Brain | Healthy | 3,233 | 10X |
| Kidney22k Multiome | Kidney | Healthy | 22,772 | 10X |
| PBMC10k Multiome | Blood | Healthy | 10,970 | 10X |

**Supplemental Figure S1.**
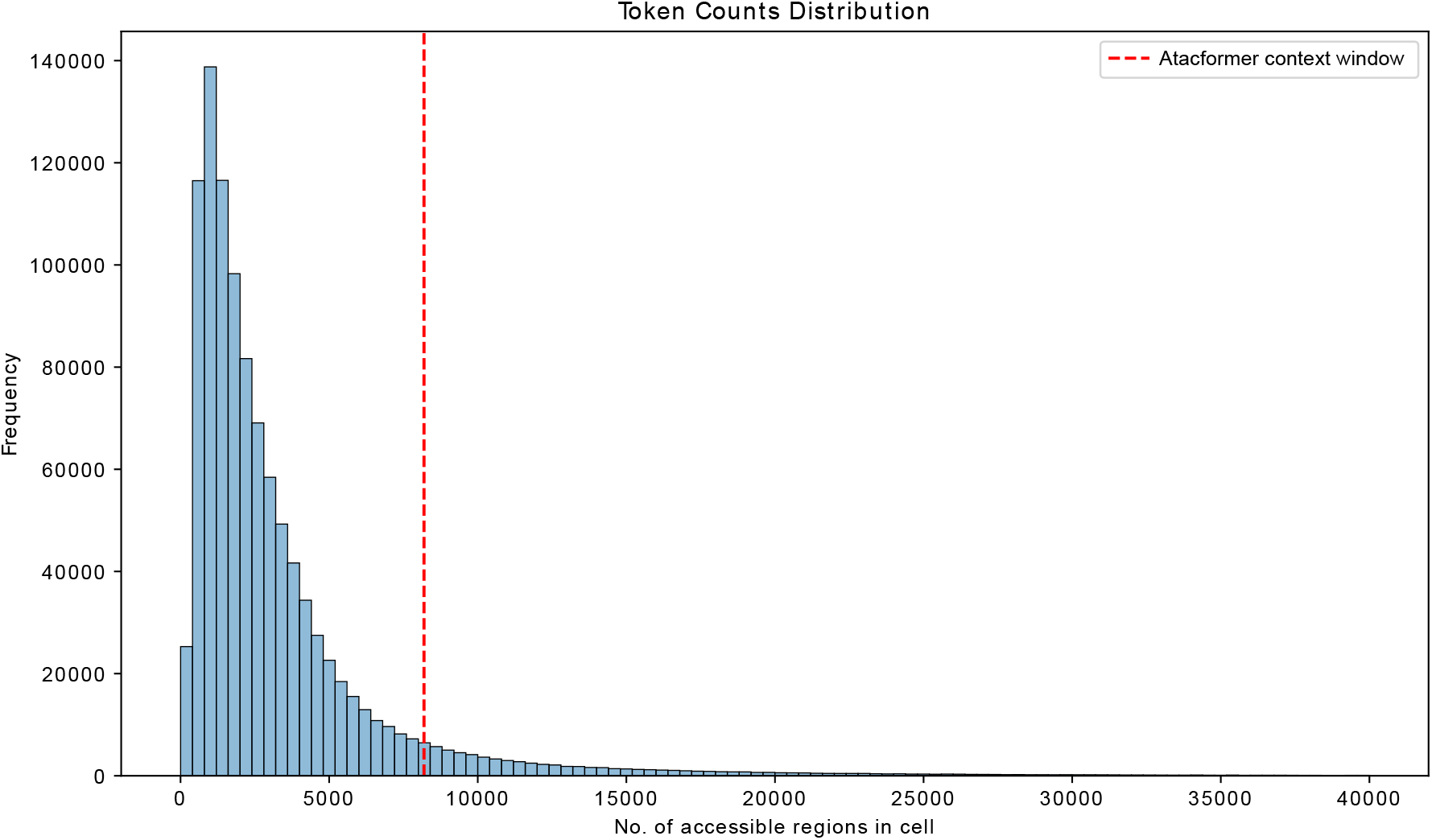
Context window distribution for all cells in the pretraining corpus. The context window is defined as the number of co-accessible regulatory elements assayed in that cell. For reference, we include the Atacformer context-window cutoff highlighting how the majority of cells are within this context window.

**Supplemental Figure S2.**
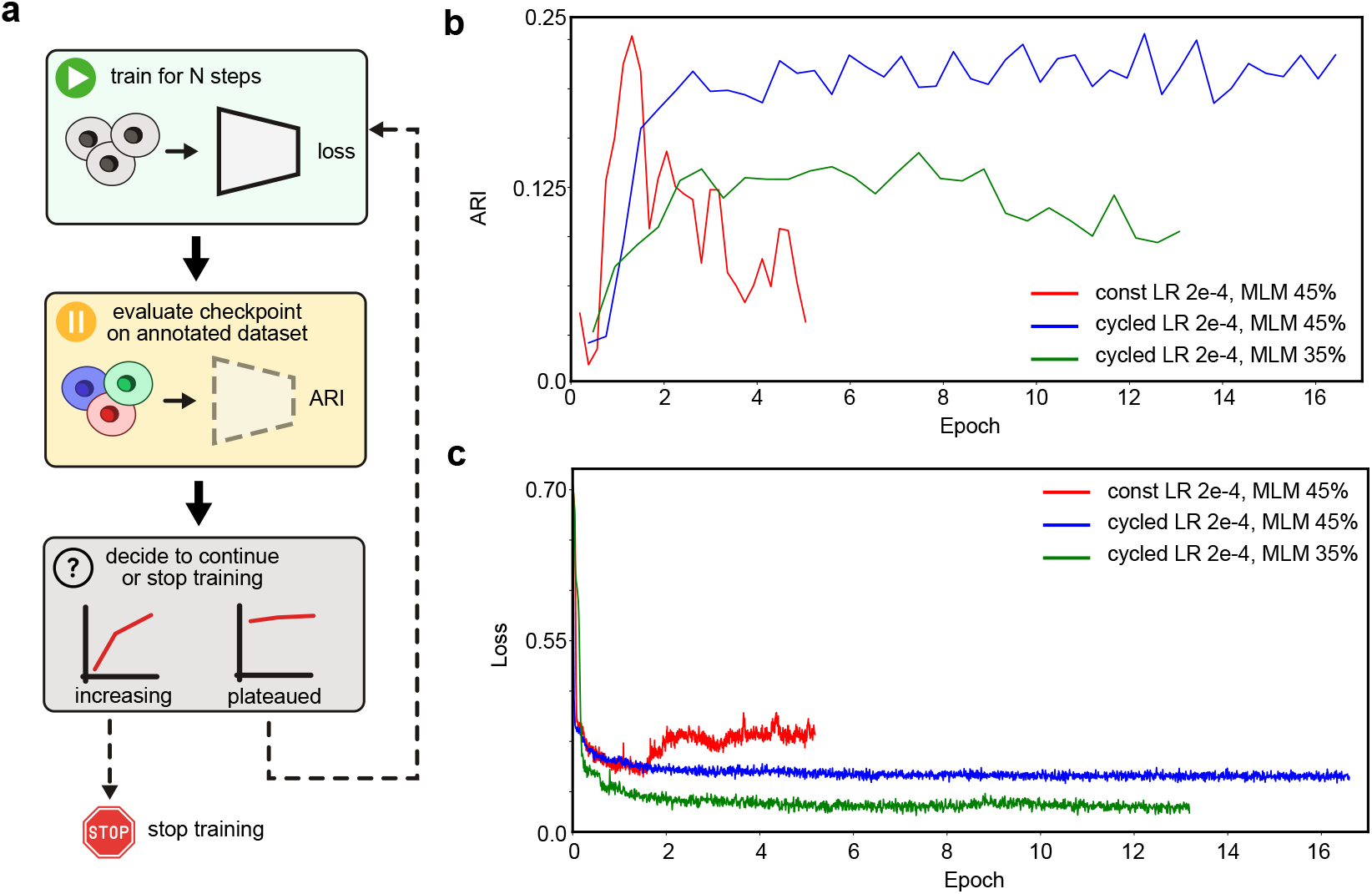
**a**. Schematic overview of Atacformer pre-training with monitored callbacks. We use these callbacks to determine whether to continue or stop training. **b**. Selected ARI callback curves for different learning rate schedules, MLM rates, and learning rates. **c**. Selected loss curves for different learning rate schedules, MLM rates, and learning rates. Notably, when the learning rate remains too high, the training becomes unstable, which is reflected in the loss curves, followed by a subsequent collapse in the ARI performance.

**Supplemental Figure S3.**
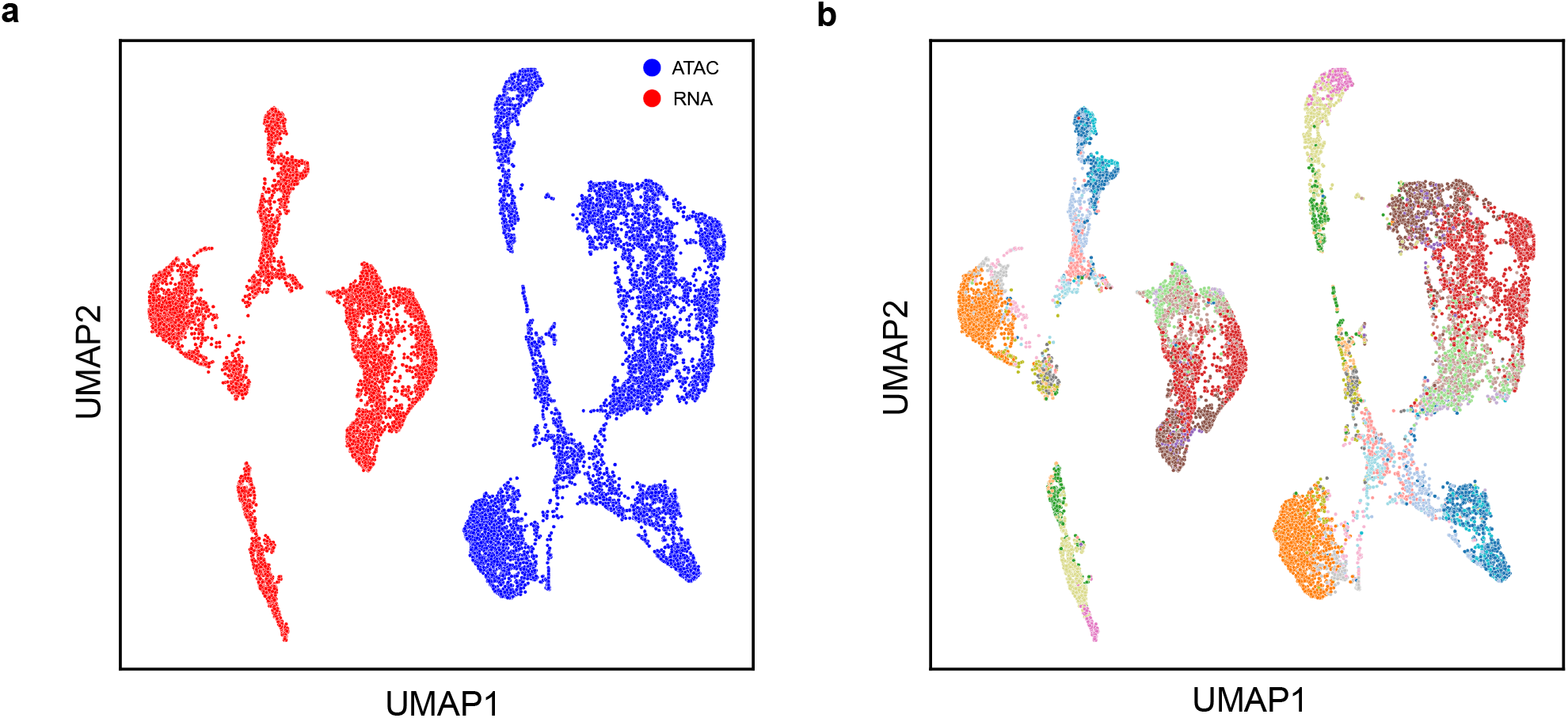
Dual UMAP visualization of both the ATAC and RNA co-embeddings. **a**. ATAC and RNA co-embeddings visualized in a shared UMAP space, colored by modality. The two modalities are divided along a shared axis. **b**. ATAC and RNA co-embeddings visualized in a shared UMAP space, colored by cell-type. Cell-type information is preserved.

**Supplemental Figure S4.**
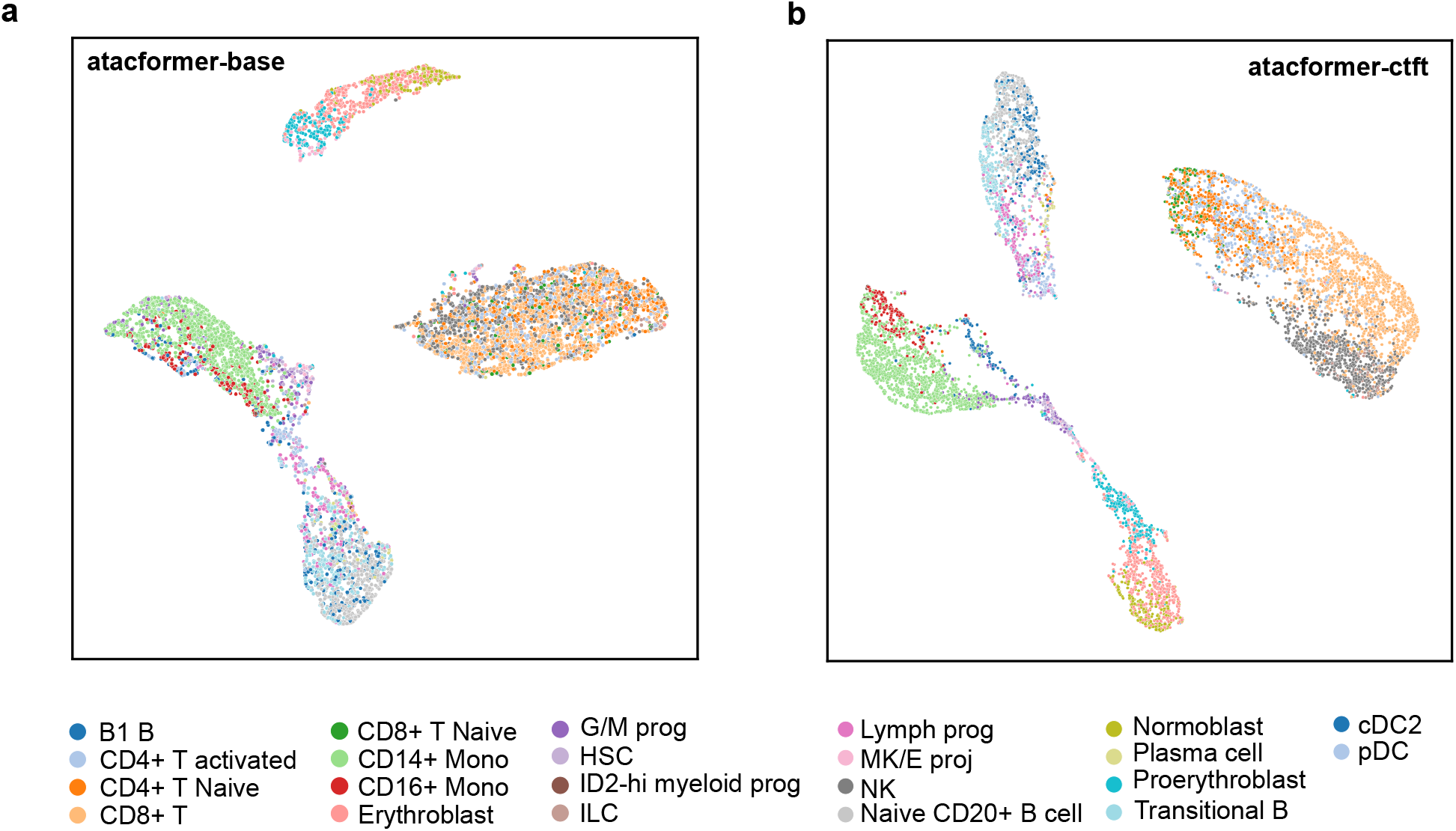
Fine-tuning Atacformer for a cell-clustering task improves latent space separation of individual cells. **a**. UMAP visualization of Luecken2021 dataset clustered using `atacformer-base` (before fine-tuning). **b**. UMAP visualization of Luecken2021 dataset clustered using `atacformer-ctft` showing a marked improvement in clustering ability.

**Supplemental Figure S5.**
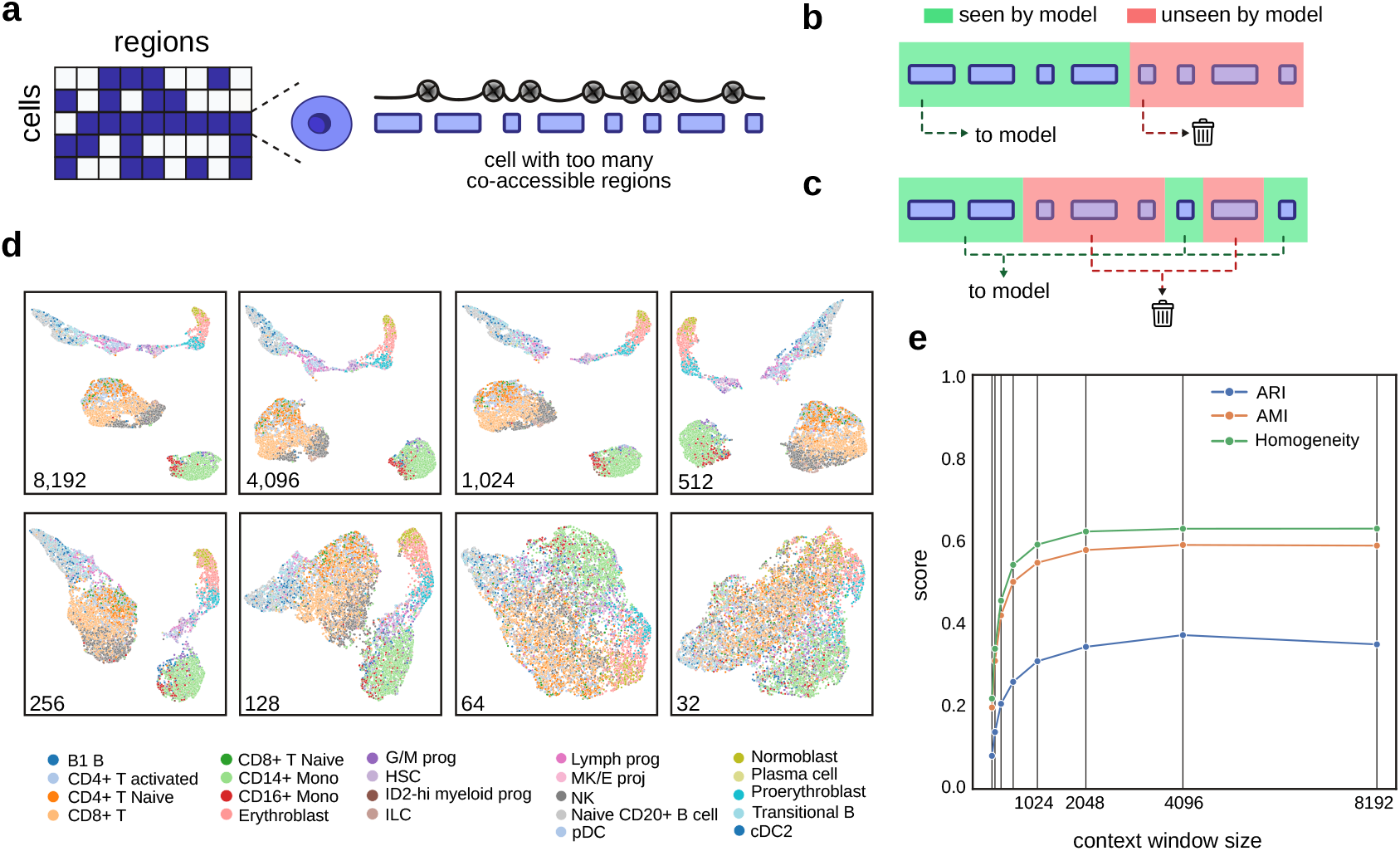
Atacformer is robust to severe degradation in context-window size. **a**. Schematic showing how cells are tokenized in the Atacformer framework. When the number of tokens in a cell exceeds the context window of the model, we must choose which tokens to drop before processing. **b**. Schematic of the cut-off method, where we simply keep the first C tokens in a cell, while disregarding the rest (C being the context-window size). **c**. Schematic of the random sample method, where we randomly sample C tokens from the cell, while discarding the rest. **d**. UMAP visualizations of embeddings generated from the Luecken2021 dataset using various context window sizes at inference time. A marked decrease in visually distinct clusters occurs after 512. **e**. Line plot of three clustering metrics as a function of context-window size. All plots and metrics utilized the ATAC encoder of the `craft-100k-hg38` model described in 2.

**Supplemental Figure S6.**
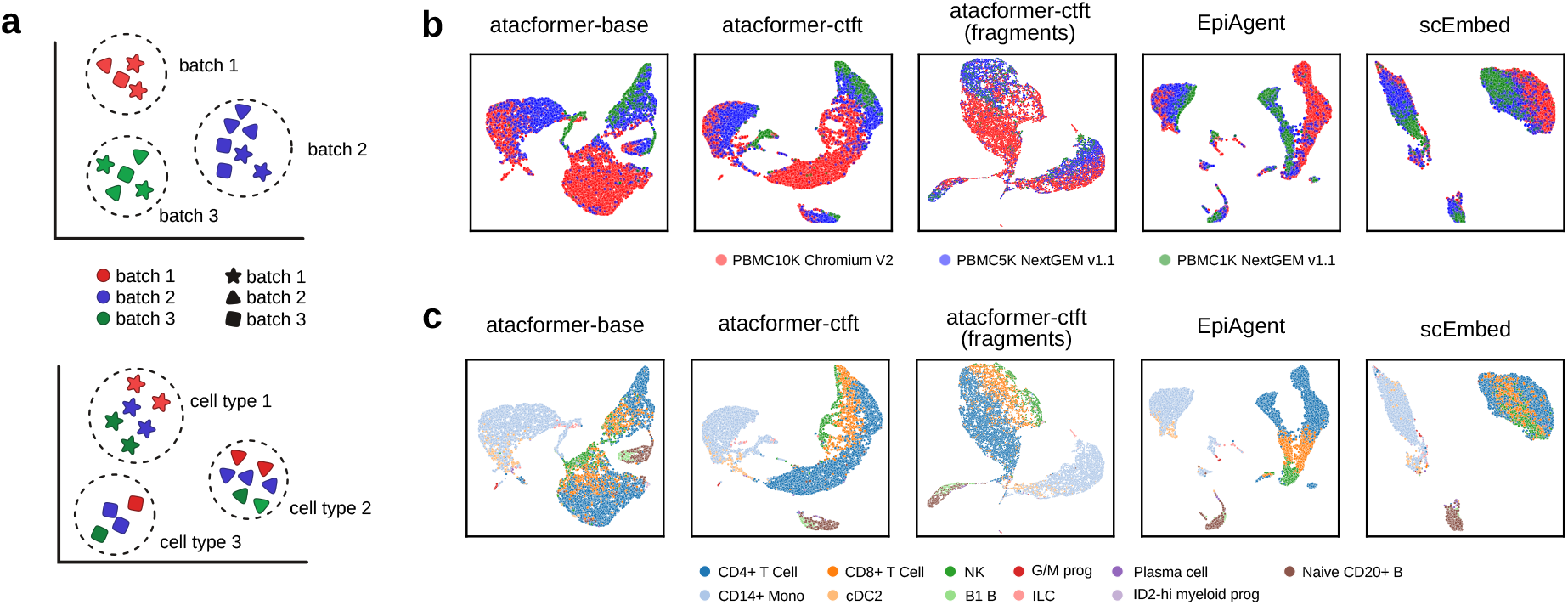
Atacformer performs strong zero-shot batch correction on processed and unprocessed data. **a**. Schematic overview of batch effects (top) and mitigation (bottom) when analyzing multiple datasets at once. **b**. UMAP visualizations of three PBMC dataset cell embeddings, colored by dataset. Atacformer visually exhibits equal or better clustering performance when directly producing embeddings of fragment files. **c**. UMAP visualizations of three PBMC dataset cell embeddings, colored by cell-type. Atacformer retains key biological information when directly producing embeddings of fragment files.

**Supplemental Figure S7.**
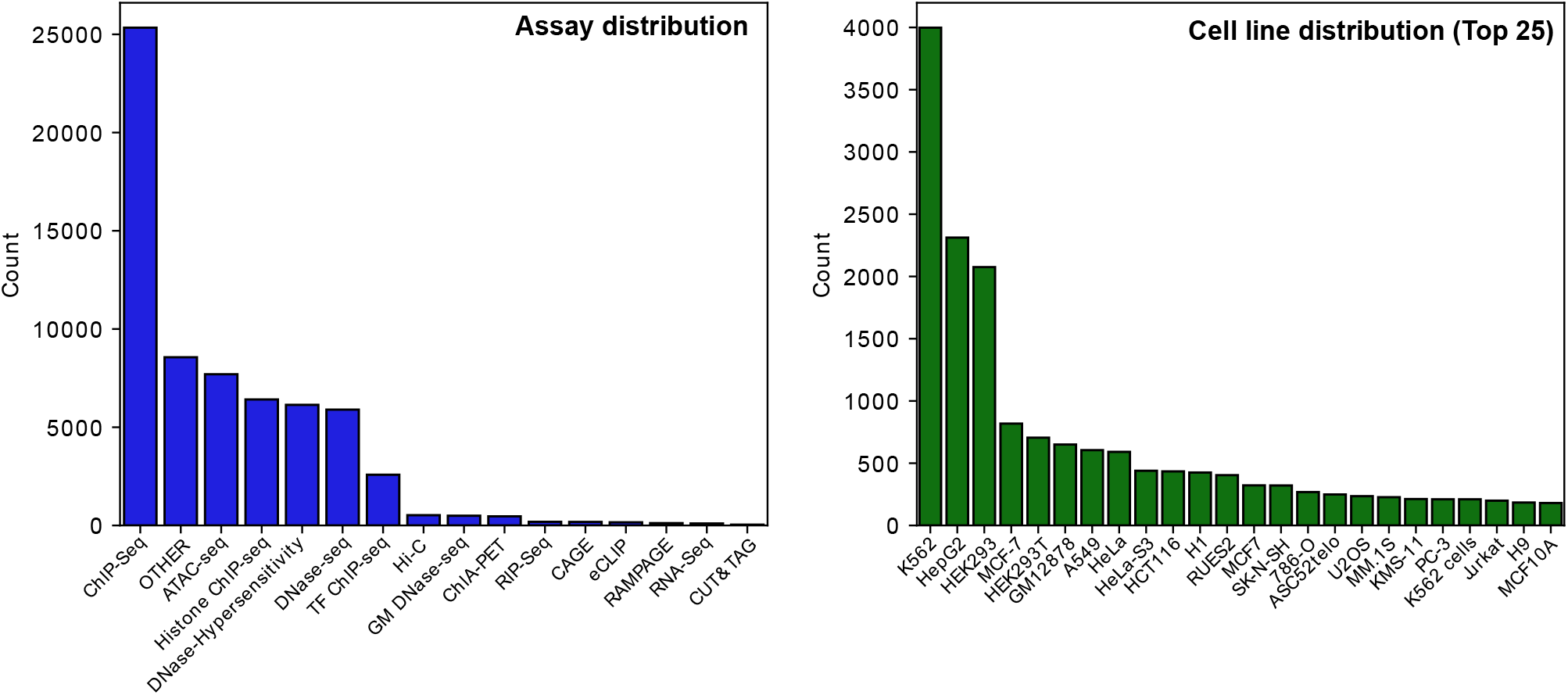
Training dataset assay and cell line distribution for the bulk-ATAC model. **a**. Distribution of assay types in the BEDbase bulk data training set. **b**. Distribution of the top 25 cell lines represented in the BEDbase bulk data training set.

**Supplemental Figure S8.**
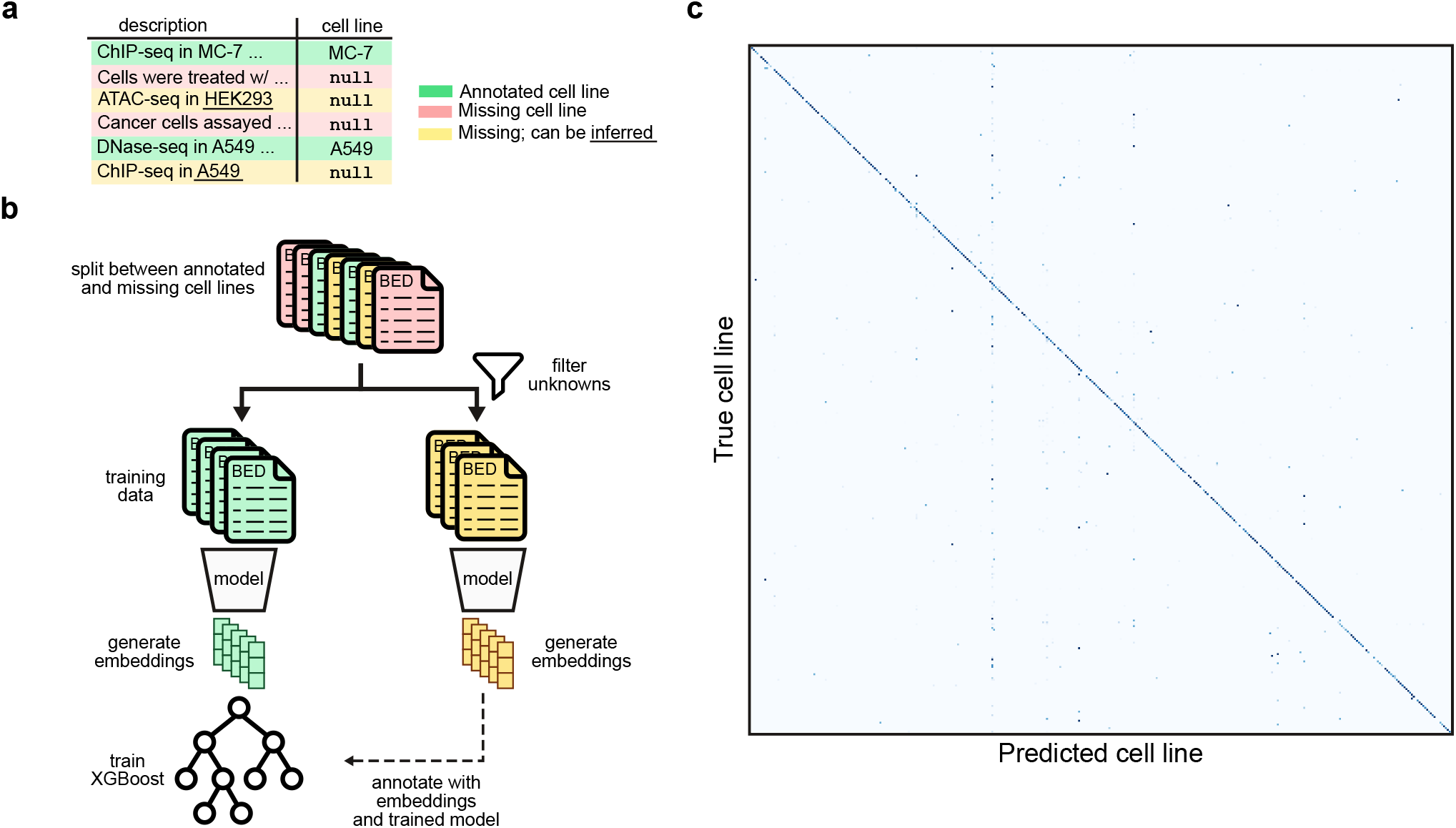
Cell line imputation for missing BEDbase data using a fine-tuned Atacformer model on bulk-ATAC data. **a**. Table demonstrating 3 different types of data rows: properly annotated rows, rows with missing cell-line annotation; and rows with missing but inferrable cell-line annotation. **b**. Schematic of the imputation procedure. **c**. Confusion matrix for entire cell line dataset showing broad agreement.

**Supplemental Figure S9.**
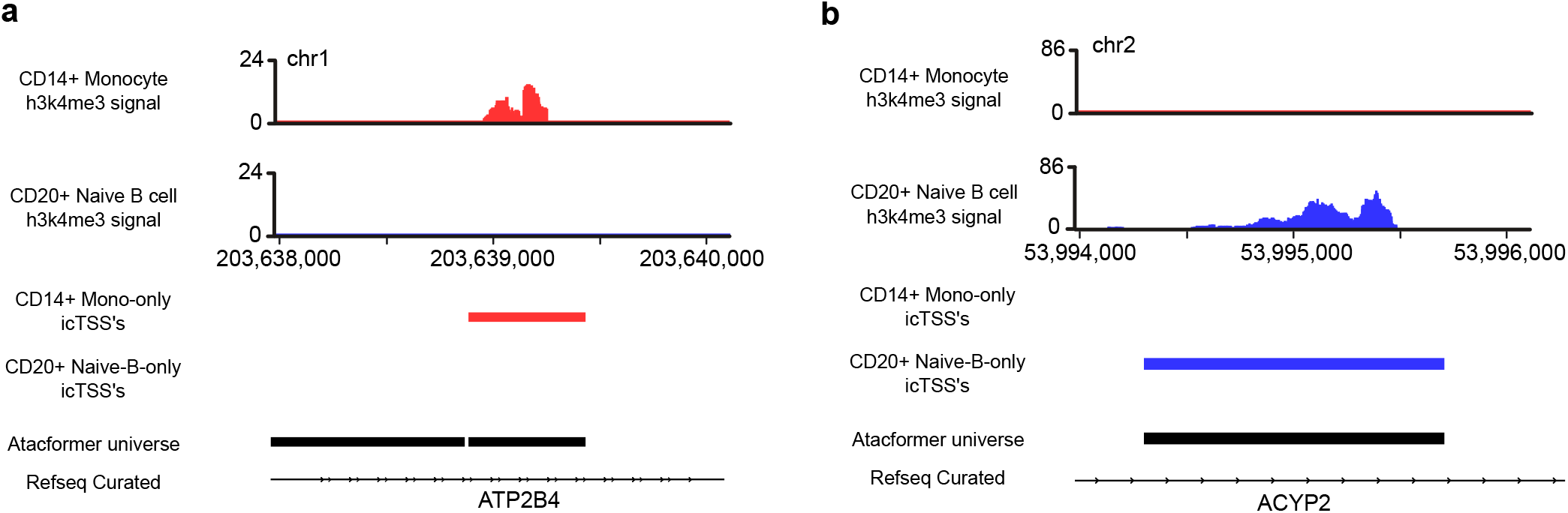
Examples of H3K4me3 enrichment in icTSS regions. **a**. Example monocyte-specific icTSS region showing H3K4me3 specifically in Monocytes. **b**. Example B-cell-specific icTSS showing H3K4me3 enrichment specifically in B cells.

**Supplemental Figure S10.**
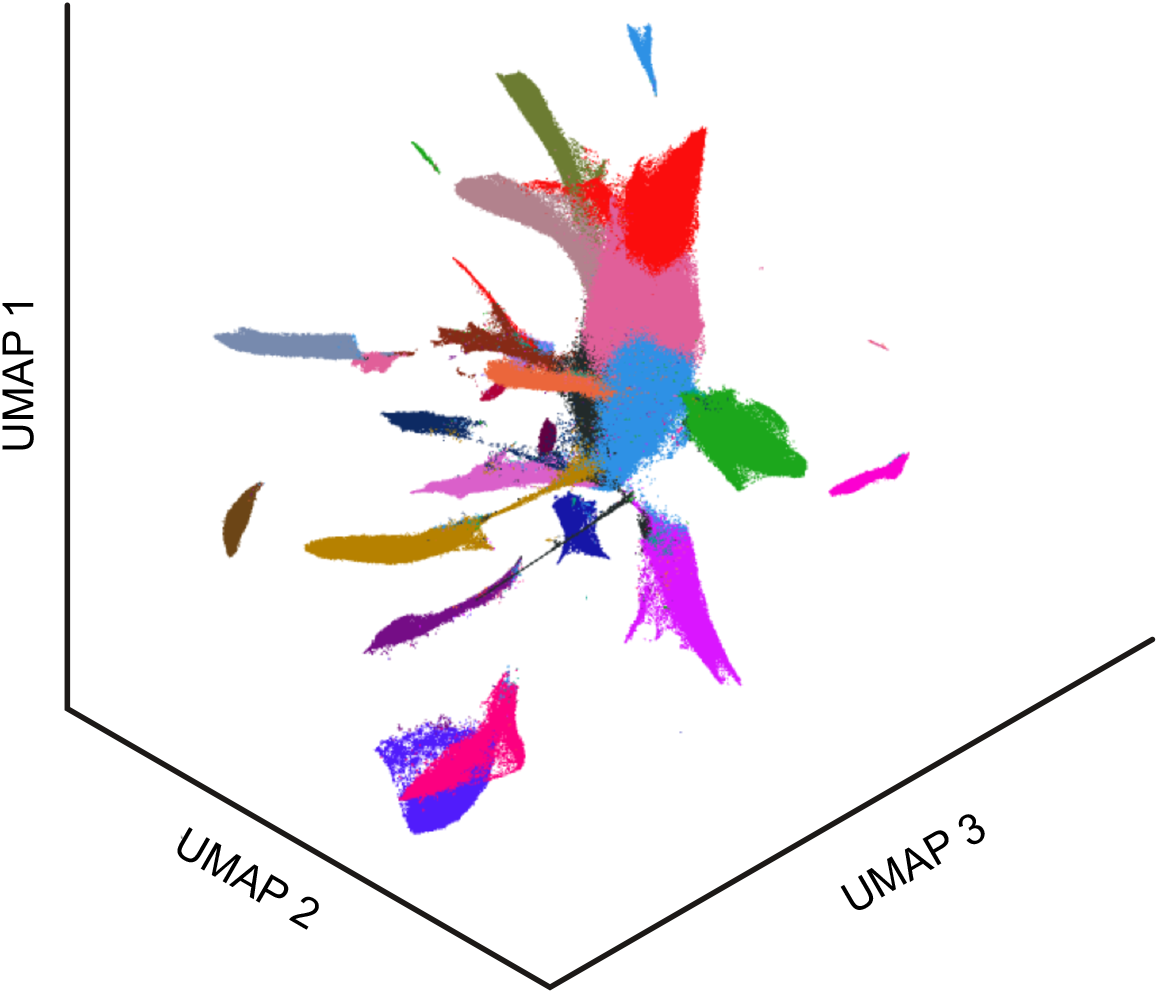
Initial clustering results of the single-cell atlas from SnapATAC2. We leverage SnapATAC2’s spectral embedding methodology and cluster using Leiden clustering.

**Supplemental Figure S11.**
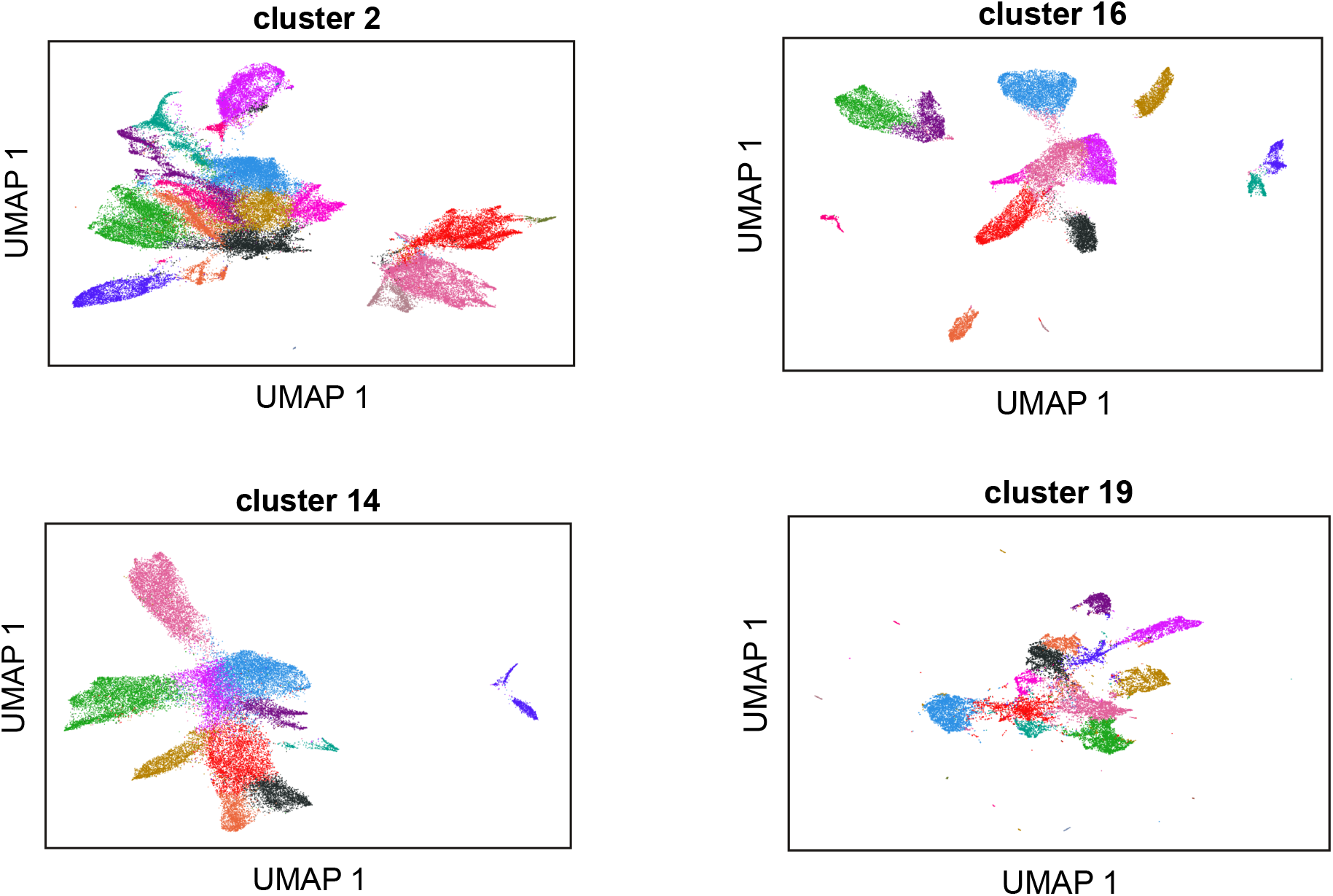
Selected sub-clustering results of the single-cell atlas from SnapATAC2. For each initial cluster, we subset the dataset and perform a secondary clustering procedure using the SnapATAC2 spectral embedding procedure. The resultant embeddings are clustered using Leiden clustering.

